# A Kinetic Scout Approach Accelerates Targeted Protein Degrader Development

**DOI:** 10.1101/2024.09.17.612508

**Authors:** Angela T. Fan, Gillian E. Gadbois, Hai-Tsang Huang, Jiewei Jiang, Logan H. Sigua, Emily R. Smith, Sitong Wu, Kara Dunne-Dombrink, Pavitra Goyal, Andrew J. Tao, William Sellers, Eric S. Fischer, Katherine A. Donovan, Fleur M. Ferguson

## Abstract

Bifunctional molecules such as targeted protein degraders induce proximity to promote gain-of-function pharmacology. These powerful approaches have gained broad traction across academia and the pharmaceutical industry, leading to an intensive focus on strategies that can accelerate their identification and optimization. We and others have previously used chemical proteomics to map degradable target space, and these datasets have been used to develop and train multiparameter models to extend degradability predictions across the proteome. In this study, we now turn our attention to develop generalizable chemistry strategies to accelerate the development of new bifunctional degraders. We implement lysine-targeted reversible-covalent chemistry to rationally tune the binding kinetics at the protein-of-interest across a set of 25 targets. We define an unbiased workflow consisting of global proteomics analysis, IP/MS of ternary complexes and the E-STUB assay, to mechanistically characterize the effects of ligand residence time on targeted protein degradation and formulate hypotheses about the rate-limiting step of degradation for each target. Our key finding is that target residence time is a major determinant of degrader activity, and this can be rapidly and rationally tuned through the synthesis of a minimal number of analogues to accelerate early degrader discovery and optimization efforts.

## Introduction

Bifunctional targeted protein degraders have revolutionized drug discovery over the past decade, providing a means to target previously undruggable functions of liganded targets, overcome inhibitor resistance mechanisms and provide enhanced selectivity over inhibition.^1^ However, targeted protein degraders can be challenging to develop, with much of the design process remaining empirical.^2^ Medicinal chemistry represents a resource-intensive component of targeted protein degrader development.^3^ Consequently, the mechanisms of targeted protein degradation, and impacts of biological and chemical variables have been intensively studied over the past 5 years.^4^

Rapid progress in understanding how biological variables contribute to a given target’s degradability in different cellular contexts has been made. We previously mapped the degradable kinome via large-scale proteomic profiling, allowing us to assign a degradability score to each degraded kinase.^5^ These data were used in combination with other proteomic, genomic and molecular modeling datasets to generate predictive machine-learning based models of degradability that can be applied proteome-wide to inform target selection.^6^ Elegant work from the Schulman lab has shown that both the activation (neddylation) state of the E3-ligase^7^, and the expression level of the hijacked E2-enzyme^8^ drive cell line and cell state dependent degradation pharmacology, informing model selection.

Chemical variables, defined as those we can change via small molecule design, have also been rigorously examined via focused experimental and theoretical studies with the aim of accelerating the optimization process and providing rational design strategies for degraders (Fig 1A).^9-15^ These models differ in their underlying structure, but agree in their predictions that the stability of the functional ternary complex has a profound impact on the expected degradation outcome. These models are used to predict how features such as the target and E3-ligase ligand binding kinetics (affinity, residence time), as well as the co-operativity (α) of the ternary complex, impact degradation activity under different biological conditions. The optimal values for each system depend on biological features such as ubiquitination and deubiquitination rate, target and E3-ligase abundance, target sequence length and proteasomal degradation rate. ^9-15^

**Figure 1.**
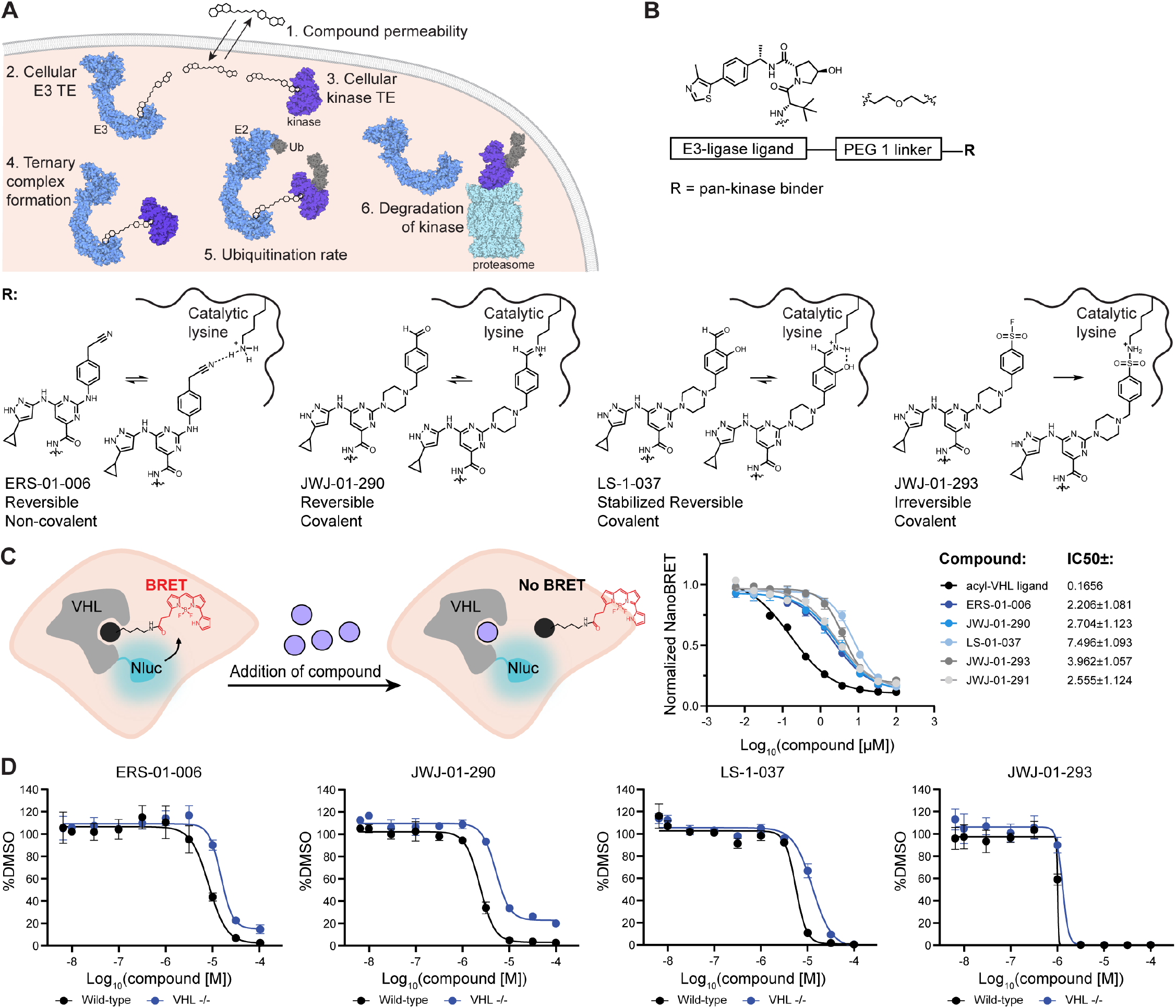
Overview of the Kinetic Scout Degrader Approach. A. Schematic showing variables that impact targeted protein degradation measured in this study. B. Design of kinetic scout degrader library. Pan-kinase binders with comparable kinome-wide selectivity profiles but distinct binding off-rates, mediated by reversible, reversible covalent and covalent interactions at the conserved lysine were employed. C. VHL NanoBRET target engagement assay to measure permeability of kinetic scout degrader library. HEK293 cells expression NanoLuc-VHL were treated with 1 µM tracer and indicated concentration of kinetic scout degrader for 2 hrs. BRET signal was normalized to DMSO BRET signal. D. Viability assay in MOLT4 and MOLT4 VHL^-/-^ cells. Cells were treated with DMSO or indicated concentration of compound for 72 hr and luminescence was measured after addition of CellTiter-Glo reagents. C.-D. Data shown as the average of *n* = 3 replicates +/-standard deviation.

Experimentally, the targeted protein degradation field has invested substantial efforts in understanding how the co-operativity of ternary complexes can be influenced by changes to the degrader linker length, composition and rigidity.^4,9^ In a typical targeted protein degrader medicinal chemistry campaign, a number of linker variants are synthesized and evaluated with the goal of influencing ternary complex formation and stability. Rational design strategies informed by structural biology of degrader-bound complexes have been successfully implemented to increase co-operativity and improve the degradation of BRD4.^9^ Recently, Ichikawa and colleagues have shown that ternary complex co-operativity of CRBN-recruiting BRD4 degraders can also be modulated by varying the component of CRBN binders oriented towards the protein-protein interface.^16^ However, the extent to which the co-operativity of a complex can be influenced by linker optimization is also determined by the features of the protein-protein interface formed in the productive ternary complex. In many cases, methodical exploration across the full range of ternary complex co-operativity values is not easily achievable for a given degrader series. Instead, structure-cooperativity relationships are empirically determined through iterative synthesis and testing.^3^

The impact of systematically fine-tuning the target residence time has also been examined, primarily in the context of the kinase target BTK, due to its relevance in cancer and the availability of matched reversible, reversible-covalent and irreversible covalent ligands targeting cysteine 481.^17-22^ These studies demonstrated that target residence time profoundly impacts degradation, and that covalent bond formation can be compatible with targeted protein degradation in some contexts. Whilst these reports support the hypothesis that fine-tuning target residence time is a viable strategy for degrader optimization, they focus on a narrow target scope, as potent cysteine-directed binders are not available for many targets-of-interest. How large of an effect fine-tuning target residence time is anticipated to have on targeted protein degraders across a wide target space remains underexplored. A rigorous understanding of how kinetic binding variables influence TPD could aid in designing more optimal initial test libraries when developing degraders for new targets, in the same way that ‘linkerology’ is now systematically explored. These data are also crucial for validation and training the next iteration of mathematical models of targeted protein degradation.^9-15^ In this study, we sought to map the impact of target residence time on ternary complex formation, ubiquitination and target protein degradation across numerous degradable targets (Figure 1A).

## Results

To systematically investigate the impact of target residence time on TPD, we required a method to vary ligand target residence time across numerous targets. Targeted protein degraders have high molecular weight, and compounds generated early in the discovery pipeline are likely to have physicochemical properties that negate efficient cellular washout, and impede accurate measurement of live-cell off-rates. To circumvent the need for large analogue libraries and live cell off-rate measurements, we leveraged a series of pan-kinase binder analogues which differ only in the functional group that interacts with the catalytic lysine (Figure 1B). Compound 3, the parent molecule, is a pan-kinase binder that interacts with the catalytic lysine via reversible hydrogen bond formation with the nitrile group.^23^ In Compound 1, a benzaldehyde participates in reversible-covalent imine formation, which is readily hydrolyzed resulting in a residence time below 10 minutes for the majority of engaged kinases, as measured by washout chemical proteomics.^24^ Stabilization of this imine by incorporation of a salicaldehyde moiety (YTP-2137, PDB: 7FIC), extends cellular residence time kinome-wide, ranging from ∼10 minutes to over 6 hours depending on the individual kinase.^24^ Finally, incorporation of a phenyl sulfonylfluoride results in irreversible covalent bond formation at the catalytic lysine kinome-wide (XO44).^25^ Importantly, each of these analogues retains a similar pan-kinome binding profile in live cells.^23-25^ We incorporated these well-characterized pan-kinome binders into VHL-recruiting targeted protein degraders, varying the regiochemistry of the lysine-targeting moiety and the linker, to form a library of compounds we termed kinetic scout degraders (Figure 1B, Figure S1). This approach allows predictable variance of the target residence time across the kinome, using a minimal set of three covalent analogues and a reversible control.

To characterize the relative cell permeability of the kinetic scout degraders we evaluated cellular VHL target engagement using a previously described dual luciferase assay where the test compound competes with dTAG^v^-1 for occupancy of VHL, resulting in a rescue of NLuc-FKBP12^F36V^ degradation (Figure S2). Promising series were re-tested in a more accurate VHL NanoBRET assay (Figure 1C).^26^ In parallel, we tested our library in cell viability assays, using isogenic parental and VHL knockout MOLT4 cells to identify compounds with degradation-dependent viability effects, observed as a reduction in cell toxicity in the VHL knockout lines (Figure 1D, Figure S3). Based on these data, we selected a set of 4 degraders that had matched regiochemistry and linker length, varying only in their lysine-targeting functional group (ERS-01-006, JWJ-01-290, LS-1-037 and JWJ-01-293 respectively). These degraders show comparable cell permeability in NanoBRET VHL engagement assays (Figure 1C) and VHL-dependent effects in viability assays (Figure 1D).

To test the impact of residence time on degradation, we performed global proteomics analysis of MOLT4 cells following treatment with ERS-01-006, JWJ-01-290, LS-1-037 or JWJ-01-293 for 5 hrs at 1 µM and 10 µM concentrations (Figure 2A). A short time point was selected to minimize the likelihood of indirect target downregulation, particularly important when profiling pan-kinase binders that profoundly change cell signaling and proteome composition at longer time points.^5^ MOLT4 cells were selected as they have high levels of active VHL, allowing for fast targeted protein degradation to occur.^7^ We observed profound differences in the degradation profiles of the kinetic scout degraders with different lysine-targeting functional groups. A total of 25 kinases were degraded by at least one compound in the series (Figure 2A). Plotting the targets that met statistical cutoffs for each molecule at one or both evaluated concentrations, we observe overlapping targets predominantly occur between compounds with closely matched reactivity at lysine, with the exception of PRKAA1 (Figure 2B). We performed orthogonal validation of the proteomics data by immunoblotting, confirming that only LS-1-037 and JWJ-01-293 degrade CDK6, at concentrations matching the proteomic profiling data (Figure 2D). As a second example, we used immunoblotting to validate that only JWJ-01-293 degraded NEK9 (Figure 2D). To ensure these effects were due to binding kinetics and not influenced by the fluoro-sulfone group in JWJ-01-293, we synthesized methyl sulfone JWJ-01-291 as a more closely matched negative control to JWJ-01-293 and confirmed no degradation. To enable cell-based studies of binary degrader-target binding, we synthesized negative control degraders where the kinase binding warhead and linker remained intact, but a diastereomer of the VHL-binding ligand was used to prevent VHL recruitment and degradation from occurring on the timescale of the assay (Figure S4A). We next used a NanoBRET cellular target engagement assay to validate that all four negative control compounds bind CDK6 in live cells with ATF-01-129, ATF-01-074, ATF-01-076 demonstrating comparable cell permeability and IC_50_ values at equilibria, and ATF-01-075 demonstrating greater potency, as expected for an irreversible compound (Figure 2E). We next performed the CDK6 NanoBRET assay to measure cellular washout kinetics, for validating that our degraders engaged with CDK6 in a comparable fashion to the parent compounds. Here, cells were treated with 10 µM degrader, washed 4 x with media, and then treated with K-10 tracer. Signal recovery was then measured over 2 hrs. The non-covalent ATF-01-129 and ATF-01-162 analogues were rapidly washed out in the CDK6 NanoBRET washout assay, indicated by a rapid return of BRET signal to baseline (DMSO). The reversible covalent (ATF-01-074, ATF-01-076) analogues demonstrated partial displacement on the timescale of the assay, consistent with a longer residence time / slower K_off_, and the irreversible covalent analogue (ATF-01-075) was not displaced consistent with covalent binding (Figure 2F, Figure S4B).

**Figure 2.**
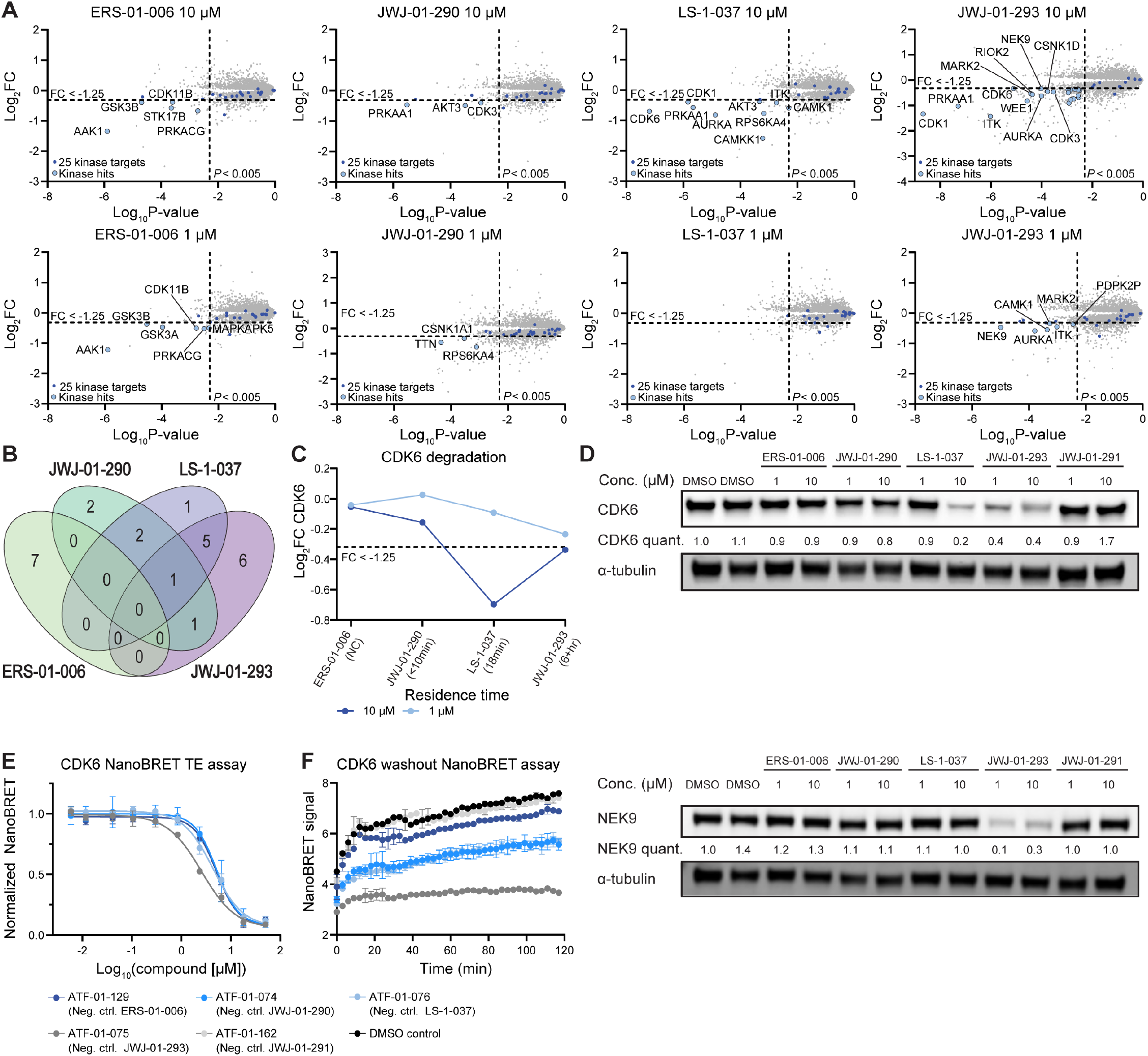
Degraded target space strongly influenced by residence time of pan-kinase recruiter. A. Global proteomics analysis of MOLT4 cells treated with the indicated compound for 5 hrs. B. Venn diagram illustrating the overlap of degraded kinases by each compound. C. Influence of kinase binder residence time on CDK6 degradation. D. Immunoblot validating compound-specific CDK6 and NEK9 degradation. MOLT4 cells were treated for 6 hrs with the indicated compounds at the indicated concentrations. Data representative of *n* = 3 biological replicates. E. CDK6 NanoBRET assay showing minimal equilibrium IC_50_ differences between reversible kinetic scout degraders. HEK293 cells expressing CDK6-NanoLuc were treated with 0.5µM K-10 tracer and indicated compound at indicated concentrations for 2 hrs. The data was background corrected by subtracting BRET signal in the absence of tracer and then normalized to DMSO BRET signal. Data shown as the average of *n* = 3 replicates +/-standard deviation. F. CDK6 NanoBRET washout assay. Cells were treated with 10 µM of the indicated compound for 2 hrs, then washed with 2 x opti-MEM+10% FBS, followed by 2 x opti-MEM. 0.5 µM K-10 tracer was added and BRET signal read over a 2 hr time course.

To examine how occupancy correlates with degradation, we next evaluated cellular target occupancy of the VHL-negative diastereomers of ERS-01-006, JWJ-01-290, LS-1-037 or JWJ-01-293 (ATF-01-129, ATF-01-074, ATF-01-076, or ATF-01-075) at equilibria across a panel of 192 kinases using NanoBRET assays in live cells at 1 µM (Figure S5). Overall, low target engagement of kinases at 1 µM degrader concentrations was observed across the series relative to control molecule XO44, consistent with reduced cell permeability of the bifunctional molecules compared to parent inhibitor controls. Expected enhancement of target engagement by irreversible molecule ATF-01-075 was observed. The CDK subfamily showed highest target occupancy by ATF-01-075, however this trend is not reflected in the degradation data. This discrepancy agrees with findings from our labs and others demonstrating that binary target occupancy does not correlate with degradation outcomes, due to the additional requirement of forming a productive ternary complex.^5^

To gain a mechanistic understanding of the drivers of differential degradation across the series, we used a sensitive VHL affinity-purification coupled to mass spectrometry method where cell lysates are supplemented with exogenous VHL-EloB-EloC to evaluate the ternary-complexes formed proteome-wide with ERS-01-006, JWJ-01-290, LS-1-037 and JWJ-01-293 (1 µM) in complex with VHL-EloB-EloC (Figure 3A). We observed enrichment of a large proportion of the kinome, reflecting the multitargeted nature of the kinase binding warheads, and indicating that the four compounds can each promote formation of ternary complexes between VHL and the majority of the kinome when VHL is not limiting (Figure 3B-C). Examining trends across the series, we observed an increase in the number and fold-change of kinase complexes enriched as the residence time of the kinase-targeting ligand increased, consistent with the predicted increase in ternary complex stability (Figure 3C).

**Figure 3.**
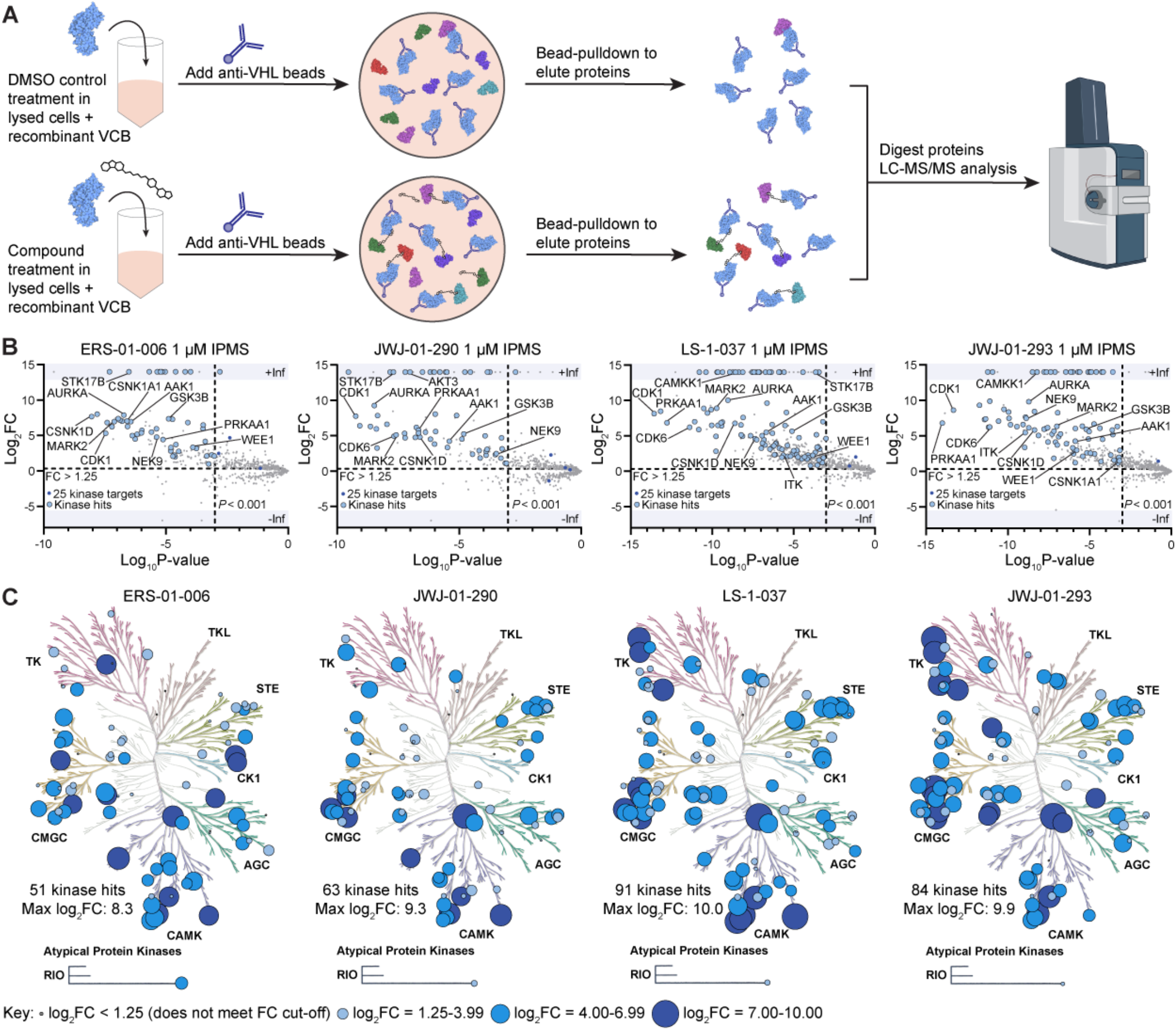
Kinetic scout degraders form ternary complexes with VHL kinome-wide. A. Schematic depicting the VHL IP/MS assay. B. Scatterplot demonstrating relative protein abundance following Flag-VHL enrichment from in-lysate treatment with 1μM of each molecule. Scatterplots display fold change in abundance to DMSO. Significant changes were assessed by moderated t-test as implemented in the limma package^78^ with log_2_FC shown on the y-axis and negative log_10_P-value on the x-axis. C. Compound-dependent VHL IP/MS enrichment of kinases plotted on a kinome tree. Illustration reproduced courtesy of Cell Signaling Technology, Inc.

To evaluate which of these complexes result in ubiquitination, we performed the E-STUB assay coupled to mass spectrometry to identify proteins ubiquitinated by VHL in the presence of 1 µM ERS-01-006, JWJ-01-290, LS-1-037 and JWJ-01-293 (Figure 4, Figure S6). To preserve the integrity of the proteome, the E-STUB assay was performed under proteasome inhibition and cells were treated with the degraders for 45 minutes followed by a 15 min biotin labeling. Consequently, not all degraded targets were enriched in the E-STUB assay. Additionally, E-STUB is validated only in easily transfectable HEK293T cells, and thus protein abundance differences between HEK293T and MOLT4 may also account for reduced target detection. Despite these caveats, we considered the E-STUB data a useful snapshot of the acute effects of the four kinetic scout degraders on targeted ubiquitination. In addition to ubiquitination of degraded targets, a number of ubiquitinated kinases were identified whose abundance did not change in our global proteomics analysis, such as CDK13. These may represent kinases that are insufficiently ubiquitinated to promote degradation, or kinases that are degraded slowly, and therefore not captured at the 5 hrs time point.

**Figure 4.**
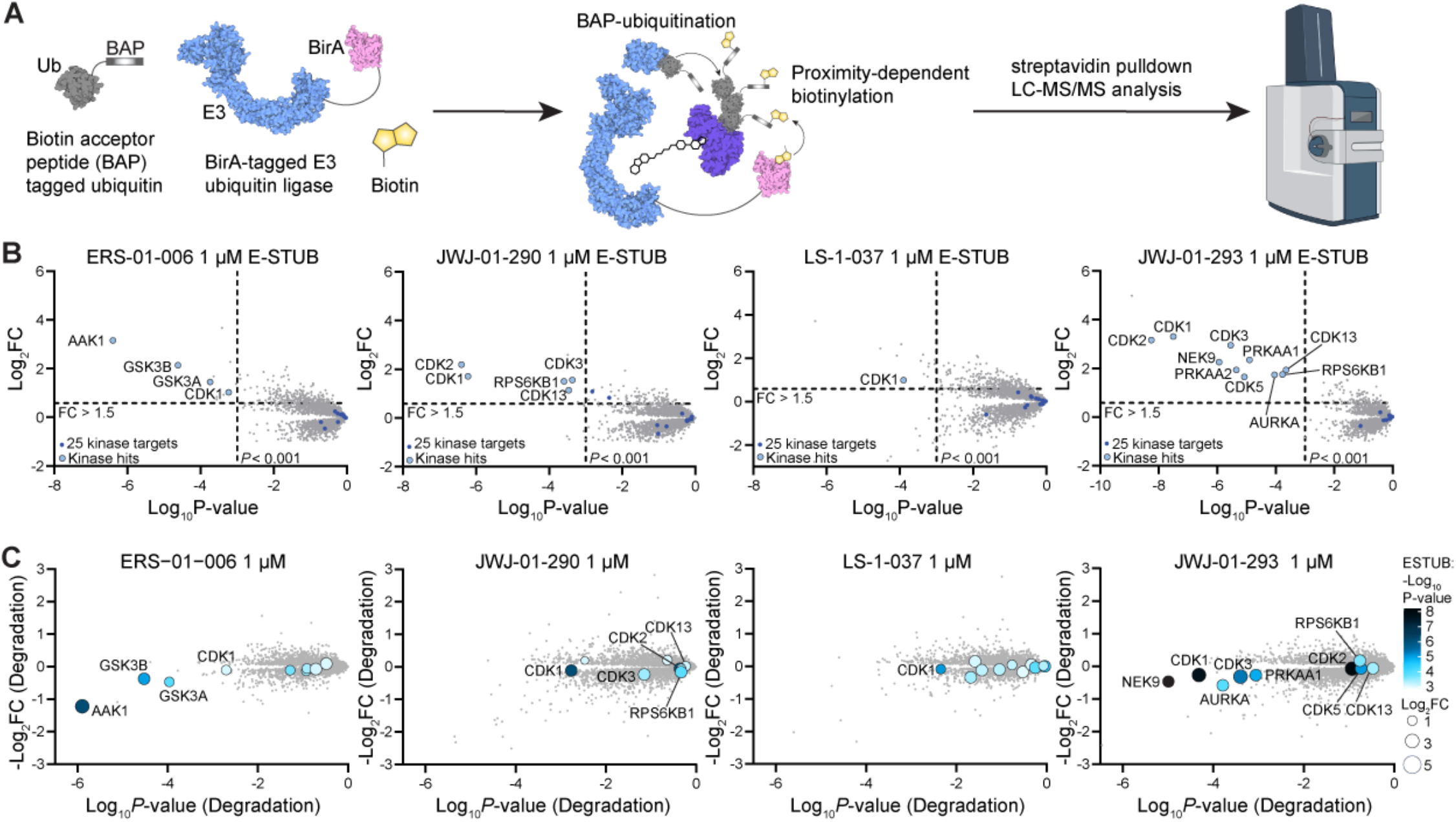
E-STUB reveals rapidly ubiquitinated kinases across the proteome. A. Schematic depicting the E-STUB assay. B. E-STUB data showing fold change in abundance of streptavidin-enriched proteins following 1 hr compound treatment in 293T VHL^-/-^ cells expressing VHL-BirA and A3-ubiquitin. C. Comparison of target-dependent degradation and ubiquitination. E-STUB data showing fold change in abundance (represented by size of dots) and statistical significance (represented by saturation of color) of B is overlaid onto a volcano plot (from Fig. 2A) of the global proteomics analysis of MOLT4 cells treated with the indicated compound for 5 hrs.

The majority of kinases complexed by VHL in the IP/MS assay were neither ubiquitinated nor degraded. We have previously shown that pan-kinase degraders can form non-productive ternary complexes that do not support degradation, which is in part supported by our ubiquitination data (Figure 4). Furthermore, unlike in our experiment, where exogenously expressed VHL was used to enhance detection of ternary complexes, in the cell, the potential kinase targets compete for a limited pool of VHL and therefore the most efficient E3-ligase:degrader:target interactions dominate the degradation pharmacology at short time points. Finally, these complexes may lead to slow ubiquitination and degradation, that occurs on a time scale beyond what we can measure with acute proteomic approaches.

To better understand how targeted protein degradation of individual kinases is impacted by differences in the kinase-binder residence time, we examined the relationship between target degradation, target occupancy, ternary complex formation, and productive ubiquitination for targets detected in multiple assays (Figure 5, Figure S7). Grouping the targets by residence-time preference allowed us to identify trends and formulate data-driven mechanistic hypotheses. Kinases such as AAK1 and STK17B were only productively ubiquitinated and degraded by fully reversible molecule ERS-01-006, but formed ternary complexes with all kinetic scout degraders, indicating for these kinases, ternary complex dissociation may be the rate limiting step for degradation. Conversely, kinases that were preferentially degraded by irreversible compound JWJ-01-293 are exemplified by AURKA, ITK and NEK9. These kinases show a greater fold-change in the enrichment of both ternary complexes and ubiquitination with longer kinase binder residence time, indicating that a greater ternary complex stability is required to achieve productive ubiquitination. CDK1 formed ternary complexes that resulted in ubiquitination with all compounds, but was preferentially degraded by JWJ-01-293, indicating that polyubiquitination may be the rate-limiting step for CDK1 degradation in this compound series. Many kinases showed profiles suggestive of both effects at work, for example CDK6 where increased enrichment of the ternary complex occurs across ERS-01-006, JWJ-01-290 and LS-1-37 resulting in degradation by LS-1-37 at 10µM, but degradation activity is reduced in the covalent analogue JWJ-01-293 at 10 µM. These results are in broad agreement with those predicted by mathematical modeling approaches, where a balance of ternary complex co-operativity and binary target residence time must be achieved for optimal degradation, and these effects vary with protein ubiquitination rate, protein abundance, and protein length. However, the residence time requirements of a given kinase for optimal degradation do not correlate with experimental or predicted scores of degradability, indicating that currently the optimal ternary complex stability for a given productive E3-ligase:degrader:target complex must be determined empirically.

**Figure 5.**
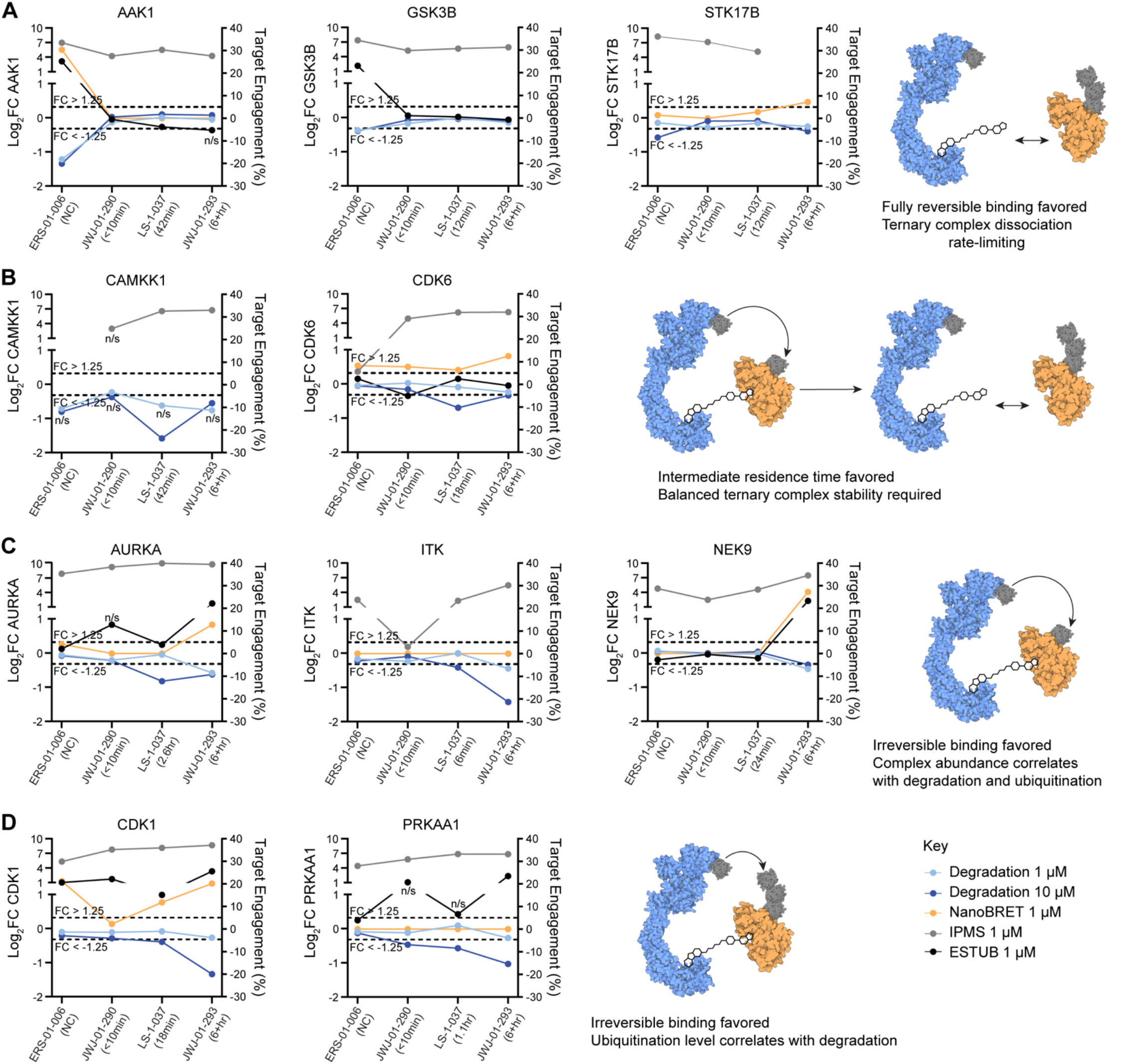
Multiparameter analysis reveals drivers of residence-time based degradation outcomes. Multiparameter profiles for representative kinases that are preferentially degraded by degraders which incorporate A. fully reversible binders. B. reversible covalent binders C.-D. irreversible covalent binders. K192 NanoBRET measurements used negative control compounds.

## Discussion

Systematic tuning of the target residence time is an under explored strategy in early targeted protein degrader development projects reported in the academic literature. Our key finding is that alterations to the off-rate of the target-binding ligand have an outsized effect on degrader efficacy and selectivity across a wide target space, comparable to varying the linker length.^5^ These findings indicate prospective degrader discovery efforts can be accelerated by incorporation of systematic ligand off-rate variance in initial degrader designs. Kinetic scout degrader libraries combined with the unbiased proteomic workflow we outline here, can identify the optimal ligand binding properties for successful degradation of a target of interest. In our study, both ternary complex formation, and the extent of ubiquitination within these complexes, were impacted by changes in ligand off-rate. Together, these data can be used to define a rational degrader optimization strategy.

We propose that the implementation of tunable reversible – covalent chemistry at lysine is an efficient method for rapid modulation of ternary complex dynamics, that can complement linkerology, when embarking on new degrader-discovery efforts. Unlike cysteine, lysine is commonly found in or adjacent to drugged enzyme active sites, and ligand binding pockets.^27^ Recent work has characterized a substantial library of lysine-targeting pharmacophores, demonstrating broad ligandability of lysines proteome-wide, indicating our approach has the potential to be applied broadly.^27^ Furthermore, although here we explore modulating ternary complex stability via systematic tuning of the target ligand residence time, a similar conceptual approach could be applied to the E3-ligase ligand.

Our previous work has shown that the creation and open sharing of unbiased chemical proteomic degradation datasets can accelerate fundamental research into degrader mechanism-of-action, by providing uniform training and test sets for machine-learning, model building and evaluation.^12^ We anticipate the datasets in this manuscript will serve as useful in training and evaluation for the mathematical modeling of targeted protein degradation and induced proximity drugs. When combined with existing models of biological degradability and *in silico* structural biology pipelines, we believe this dataset and approach will ultimately aid development of better predictive modeling with which to guide the development of bifunctional induced proximity molecules.

## Supporting information

Synthetic methods

Biology methods

## Funding

This project was supported by an NSF CAREER to F.M.F. (NSF CHE-2339705). A.T.F. was supported by an NIH Chemistry-Biology Interfaces Training Grant (T32GM146648). G.E.G. was supported by an NIH Molecular Biophysics Training Grant (T32GM139795). F.M.F. was supported by NIH (DP2NS132610). A.J.T. was supported by a UCSD Distinguished Graduate Student Award and the NIH/NCI Cancer Cell Signaling & Communication Training Grant (T32CA009523). P.G. was supported by a UCSD Undergraduate Summer Research Award. L.H.S was supported by an NIH Medical Scientist Training Grant T32GM007198-49.

## Acknowledgements

We thank Promega for providing reagents and assistance in running the kinase 192 NanoBRET profiling assay.

## Author contributions

A T F, J J, L H S, E R S, G J P, K D-D, performed small molecule synthesis. G E G, P G, A J T, performed cellular assay development and evaluation of molecules. GEG, ATF, AJT performed data analysis. ATF, GEG performed figure preparation. H-T H, performed the E-STUB assay. K A D performed proteomics experiments and data analysis. W S, E S F, F M F supervised the study, and performed funding acquisition. KAD and FMF conceived the study. FMF wrote the manuscript with input and edits from all authors.

## Conflict of Interest Statement

K.A.D receives or has received consulting fees from Kronos Bio and Neomorph Inc. H.-T.H has no conflict of interest. W.R.S. is a Board or SAB member and holds equity in Delphia Therapeutics, Ideaya Biosciences, Red Ridge Bio, Scorpion Therapeutics and has consulted for Array, Astex, CJ Biosciences, Epidarex Capital, Ipsen, Merck Pharmaceuticals, Pierre Fabre, Sanofi, Servier and Syndax Pharmaceuticals and receives research funding from Bayer Pharmaceutical, Bristol-Myers Squibb, Boehringer-Ingelheim, Ideaya Biosciences, Calico Biosciences, and Servier Pharmaceuticals. W.R.S. is a co-patent holder on EGFR mutation diagnostic patents. F.M.F. is a scientific co-founder and equity holder in Proximity Therapeutics, and was previously a scientific advisory board member (SAB) of Triana Biomedicines. F.M.F. is or was recently a consultant or received speaking honoraria from RA Capital, Eli Lilly and Co., Sorrento Pharma, Plexium Inc, Sygnature Discovery, Neomorph Inc. and Tocris BioTechne. The Ferguson lab receives or has received research funding or resources in kind from Ono Pharmaceutical Co. Ltd, Promega Corp, Eli Lilly and Co., and Merck and Co. F.M.F.’s interests have been reviewed and approved by the University of California San Diego in accordance with its conflict-of-interest policies. E.S.F. is a founder, scientific advisory board (SAB) member, and equity holder of Civetta Therapeutics, Proximity Therapeutics, Stelexis Biosciences, and Neomorph, Inc. (also board of directors). He is an equity holder and SAB member for Avilar Therapeutics, Photys Therapeutics, and Ajax Therapeutics and an equity holder in Lighthorse Therapeutics and Anvia Therapeutics. E.S.F. is a consultant to Novartis, EcoR1 capital, Odyssey and Deerfield. The Fischer lab receives or has received research funding from Deerfield, Novartis, Ajax, Interline, Bayer and Astellas.

---

**Supporting Figure 1.**
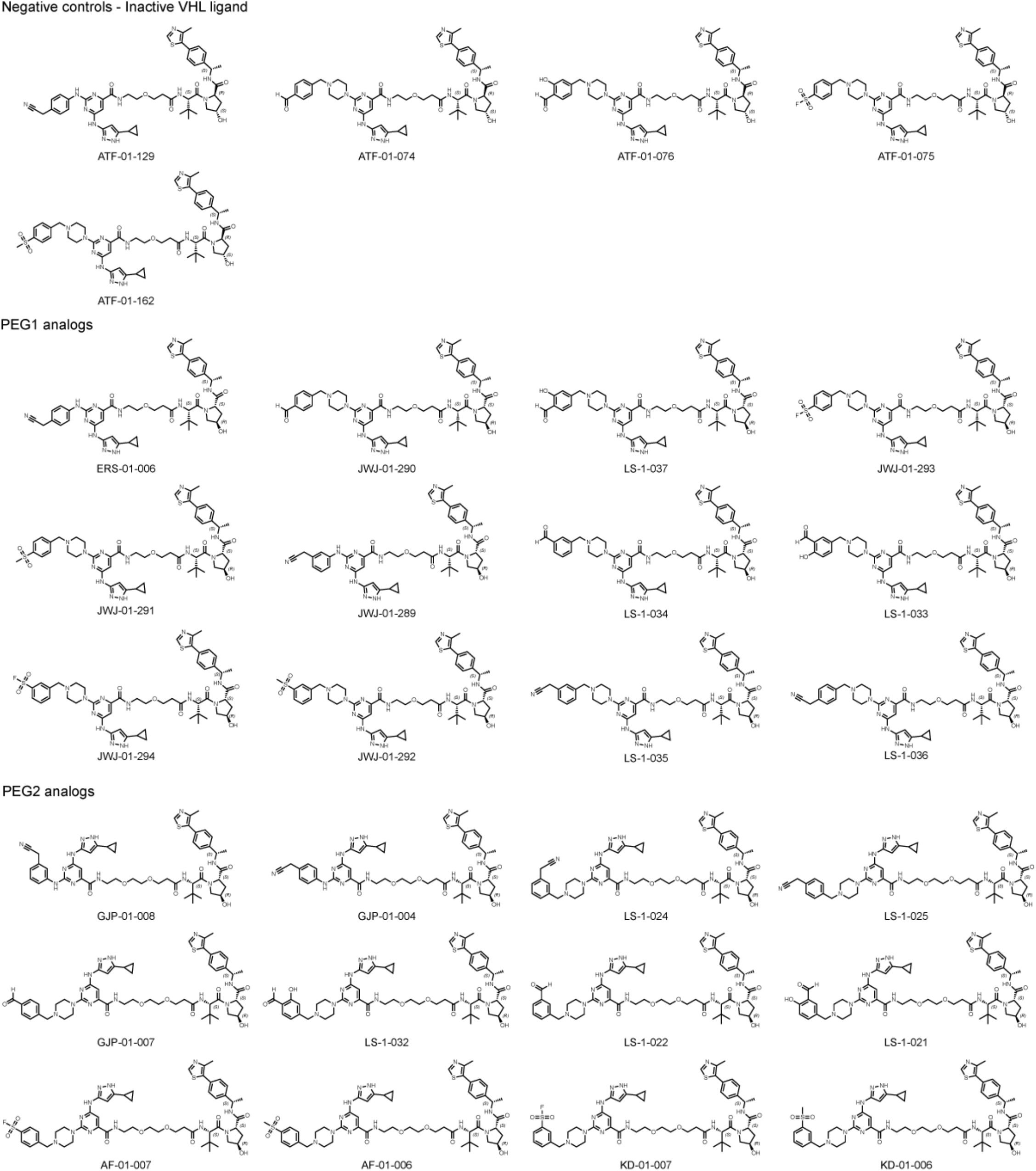
Chemical Structures of Kinetic Scout Degrader Library Compounds.

**Supporting Figure 2.**
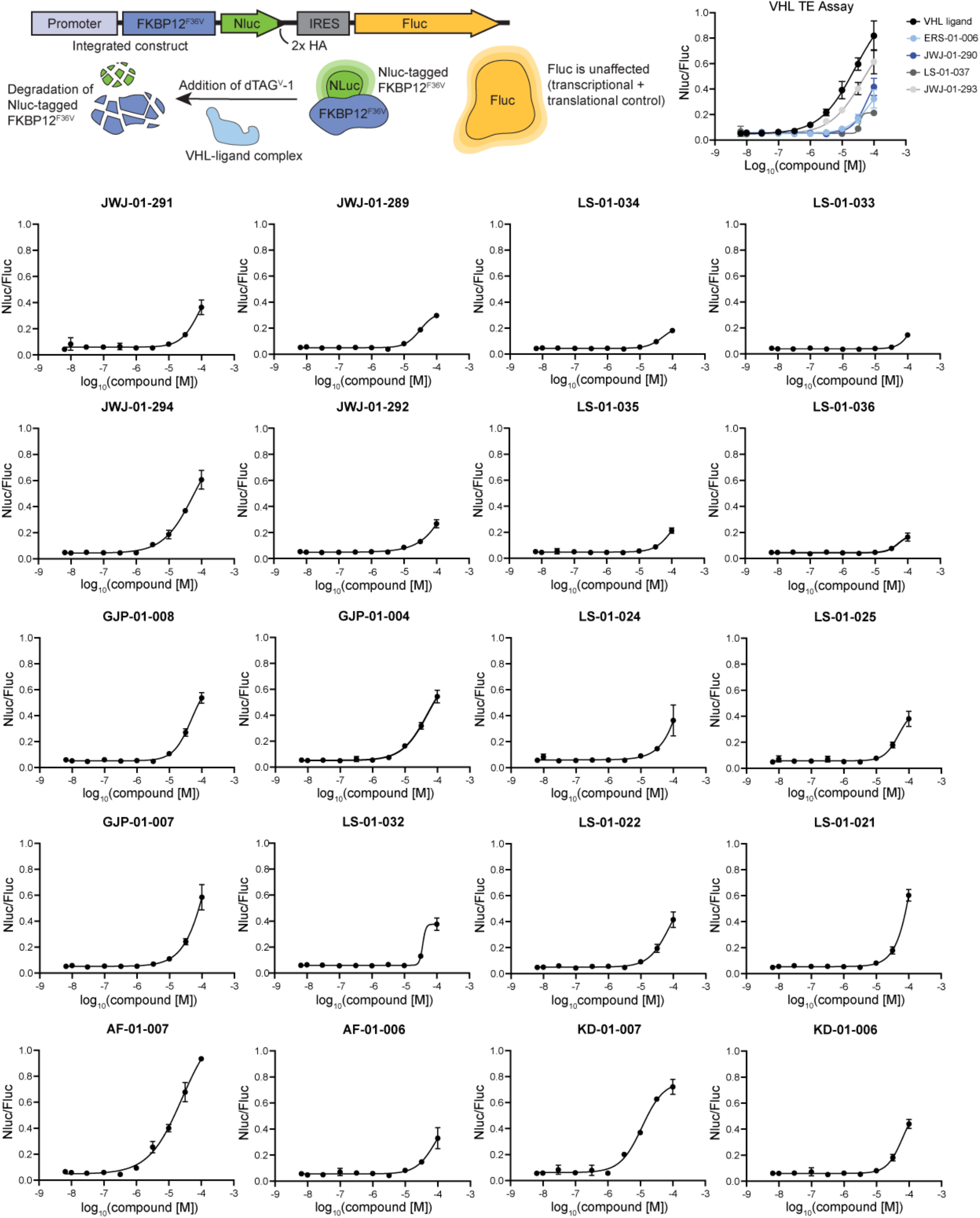
Cellular VHL Engagement of Kinetic Scout Degrader Library. VHL cellular target engagement assay measures competitive displacement of dTAG^v^-1 in cells. HEK293FT-FKBP12^F36V^-NanoLuc cells were treated with 100 nM dTAG^v^-1 and indicated concentration of compound for 6 hr and luminescence was measured using a ClarioSTAR Plus microplate reader. Data shown as the average +/-S.D. of *n* = 3 replicates.

**Supporting Figure 3.**
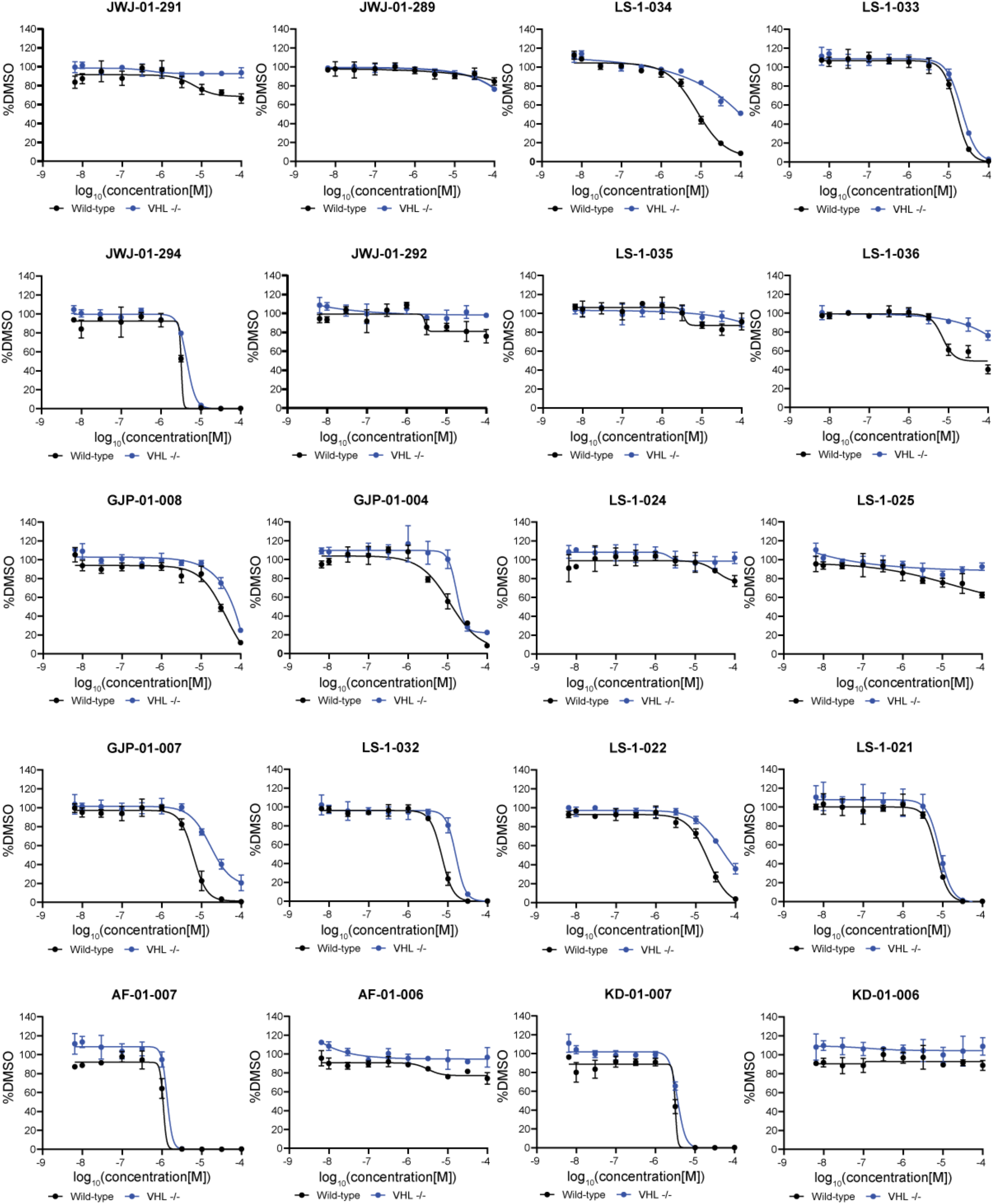
VHL-dependent Cell Viability Effects of Kinetic Scout Degraders. Viability assay in MOLT4 and MOLT4 VHL^-/-^ cells. Cells were treated with DMSO or indicated concentration of compound for 72 hr and luminescence was measured after addition of CellTiter-Glo reagents. Data shown as the average +/-S.D. of *n* = 3 replicates.

**Supporting Figure 4.**
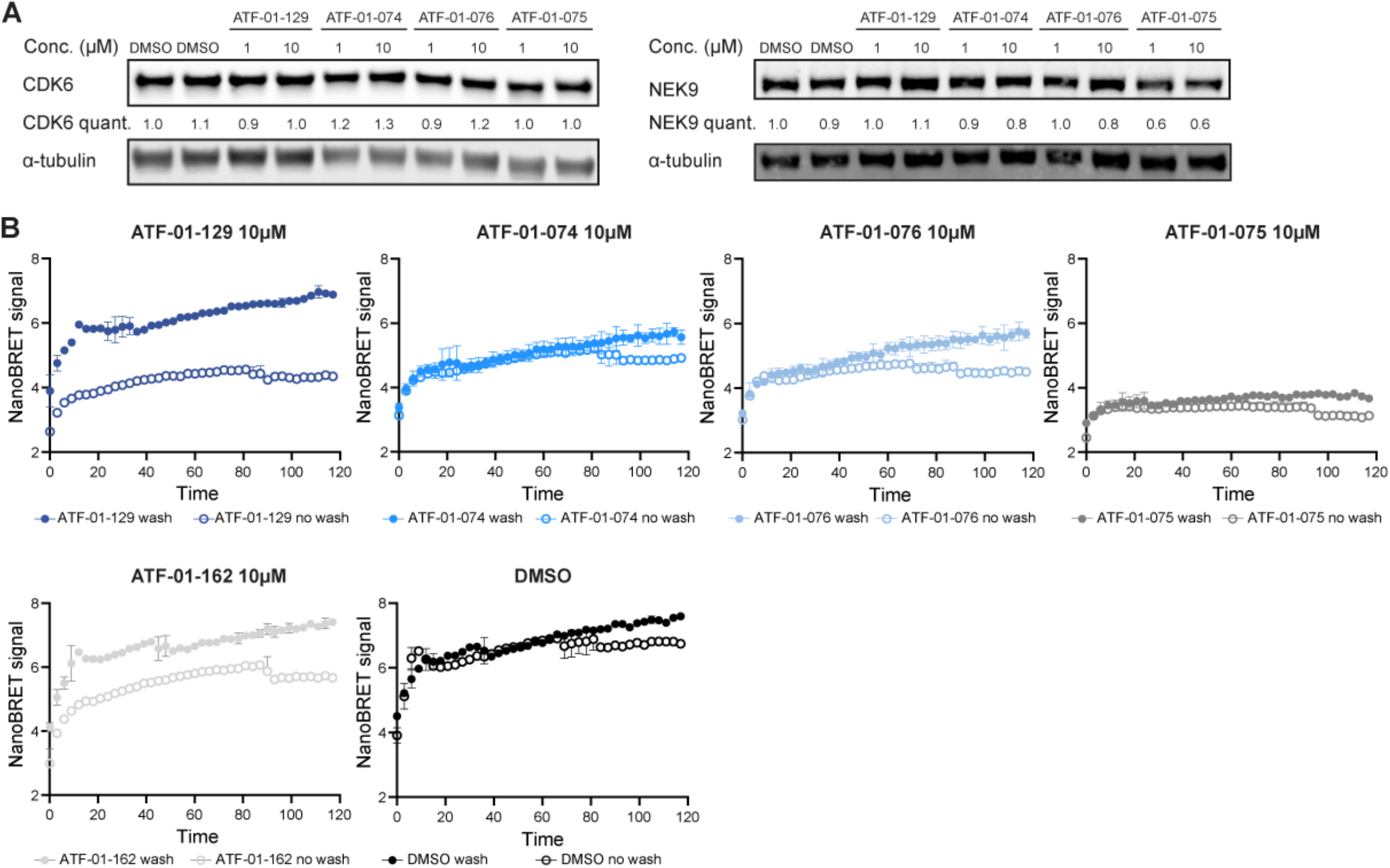
CDK6 NanoBRET washout experiment. A. Immunoblot validating no degradation of CDK6 and NEK9 with VHL inactive compounds. MOLT4 cells were treated for 6 hrs with the indicated compounds at the indicated concentrations. B. Cells expressing CDK6-NanoLuc construct were treated with DMSO or indicated compound for 2 hr and then media was replaced twice with Opti-MEM + 10% FBS and then twice with Opti-MEM for wells intended to be washed. 0.5µM K-10 tracer and Complete Substrate Plus Inhibitor Solution was added then NanoBRET signal was measured. Data shown as the average +/-S.D. of n= 2 replicates.

**Supporting Figure 5.**
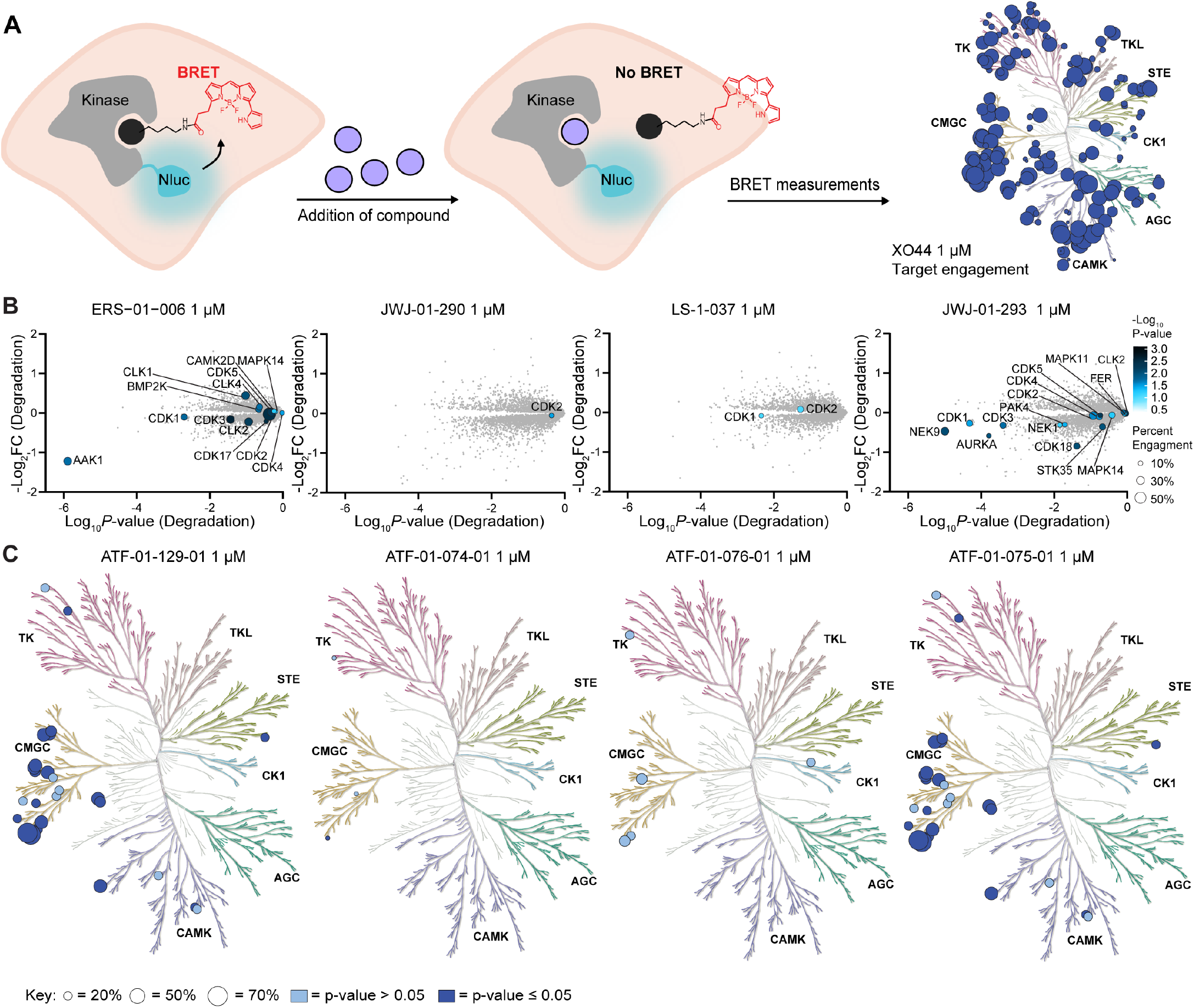
Equilibrium kinase target engagement is not a driver of degradation. A. Schematic depicting the NanoBRET K192 profiling assay. B.-C. Live cell K192 NanoBRET assay used to determine percent engagement of kinases. HEK293 cells expressing kinase-NanoLuc were treated with K-10 tracer and 1 µM compound. The percent engagement is the percent reduction of BRET signal after compound treatment compared to DMSO. Data shown as the average of *n* = 3 replicates. *P*-value was calculated using Student’s t-test. B. Comparison of cellular target engagement and degradation. Proteomic data as in Fig. 2. Percent engagement data showing compound binding (represented by size of dots) and statistical significance (represented by saturation of color) of K192 NanoBRET assay is overlaid onto a volcano plot (from Fig. 2A) of the global proteomics analysis of MOLT4 cells treated with the indicated compound for 5 hrs. Percent engagement data was measured using negative control compounds. C. Percent target engagement plotted on KinMap. Illustration reproduced courtesy of Cell Signaling Technology, Inc.

**Supporting Figure 6.**
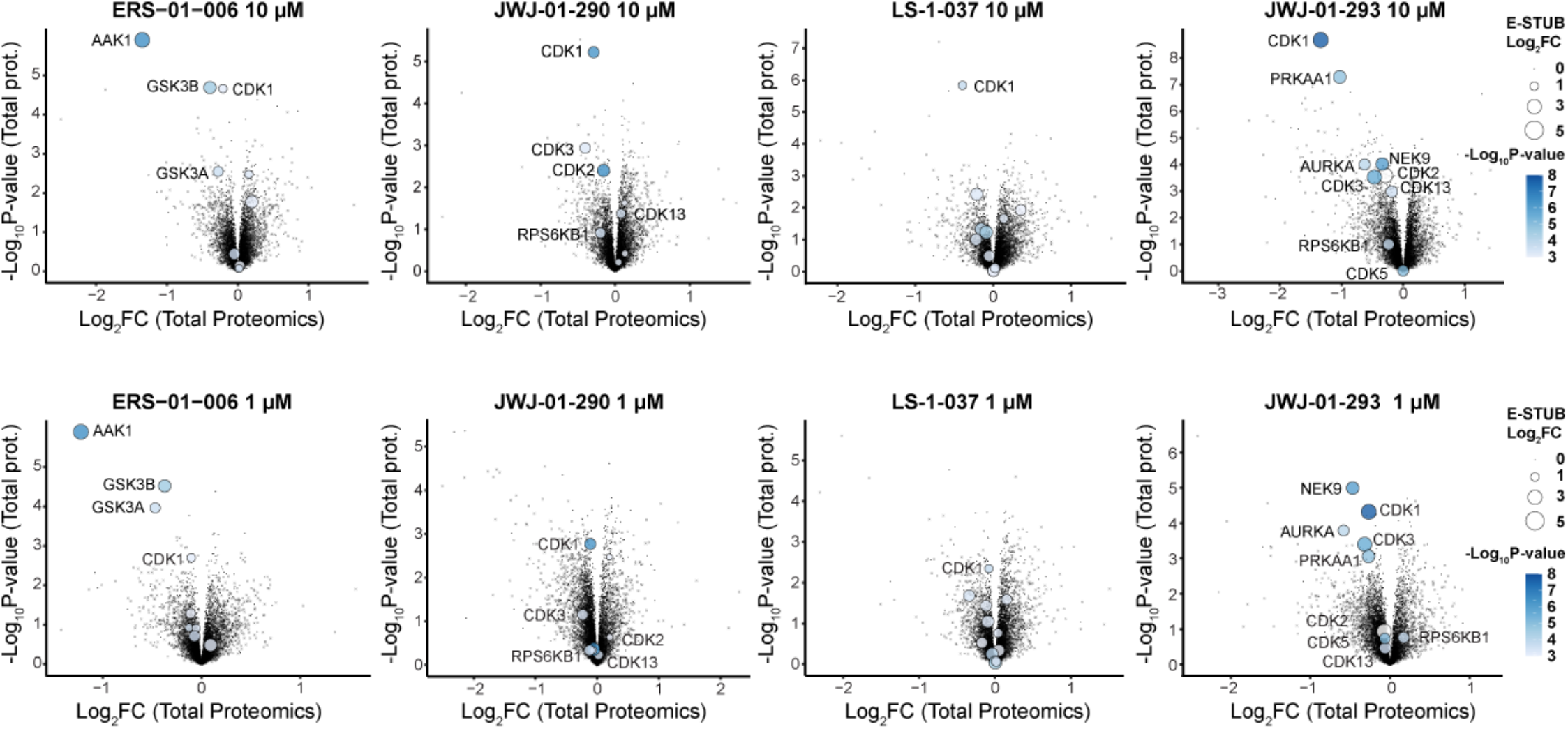
Comparison of degraded proteins and their enrichment in the E-STUB assay. E-STUB data showing fold change in abundance (represented by size of dots) and statistical significance (represented by saturation of color) of streptavidin-enriched proteins following 1 hr compound treatment in 293T VHL^-/-^ cells expressing VHL-BirA and A3-ubiquitin. The data is overlaid onto a volcano plot (from Fig. 2A) of the global proteomics analysis of MOLT4 cells treated with the indicated compound for 5 hrs. Proteins identified in global proteomics that are not detected by E-STUB are shown as a cross.

**Supporting Figure 7.**
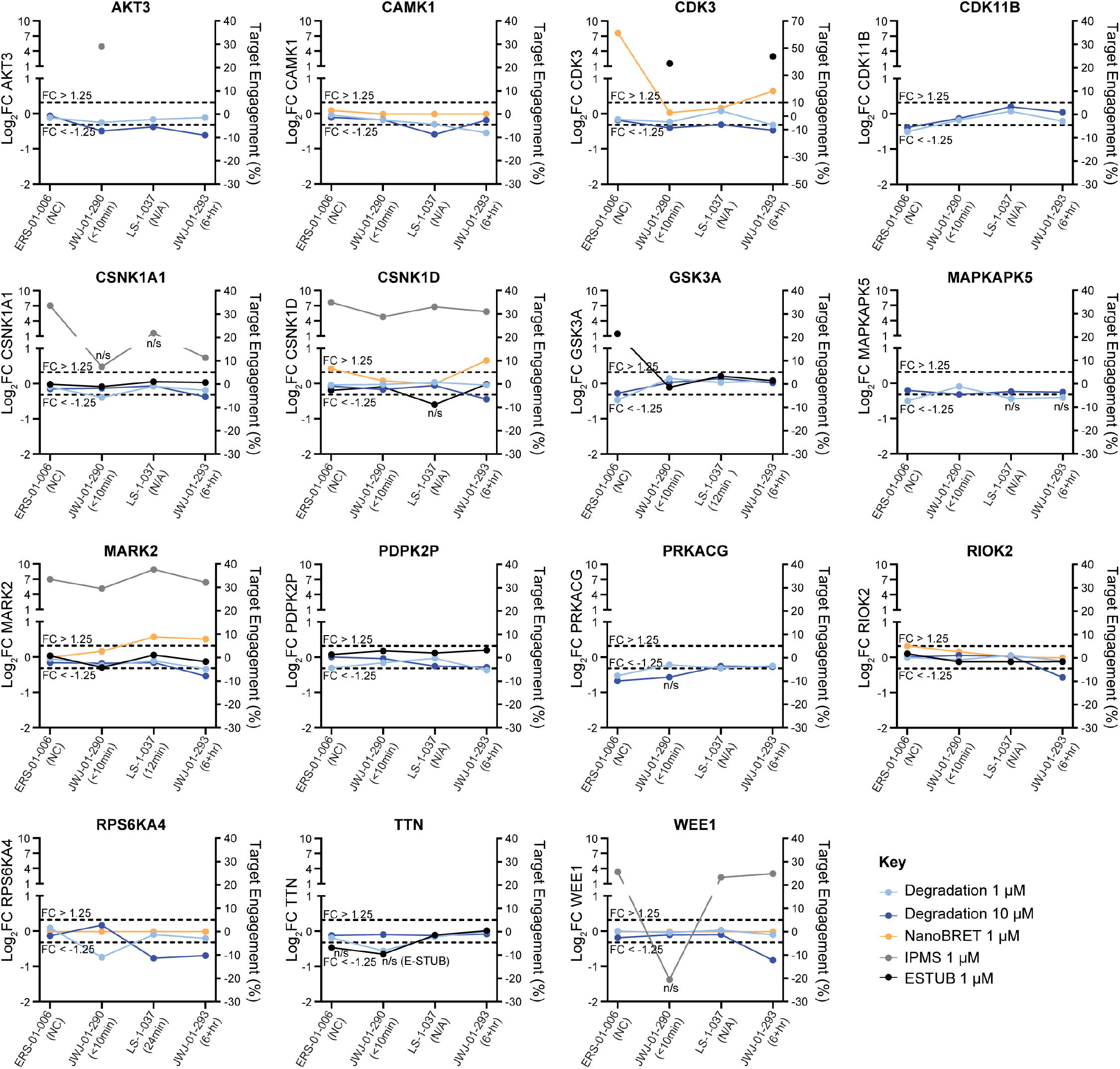
Multiparameter analysis reveals drivers of residence-time based degradation outcomes for additional degraded kinases. Multiparameter profiles for kinases not shown in Figure 5. K192 NanoBRET measurements used negative control compounds.

