## Supplementary material for "A Kinetic Scout Approach Accelerates Targeted Protein Degrader Development": Synthetic methods

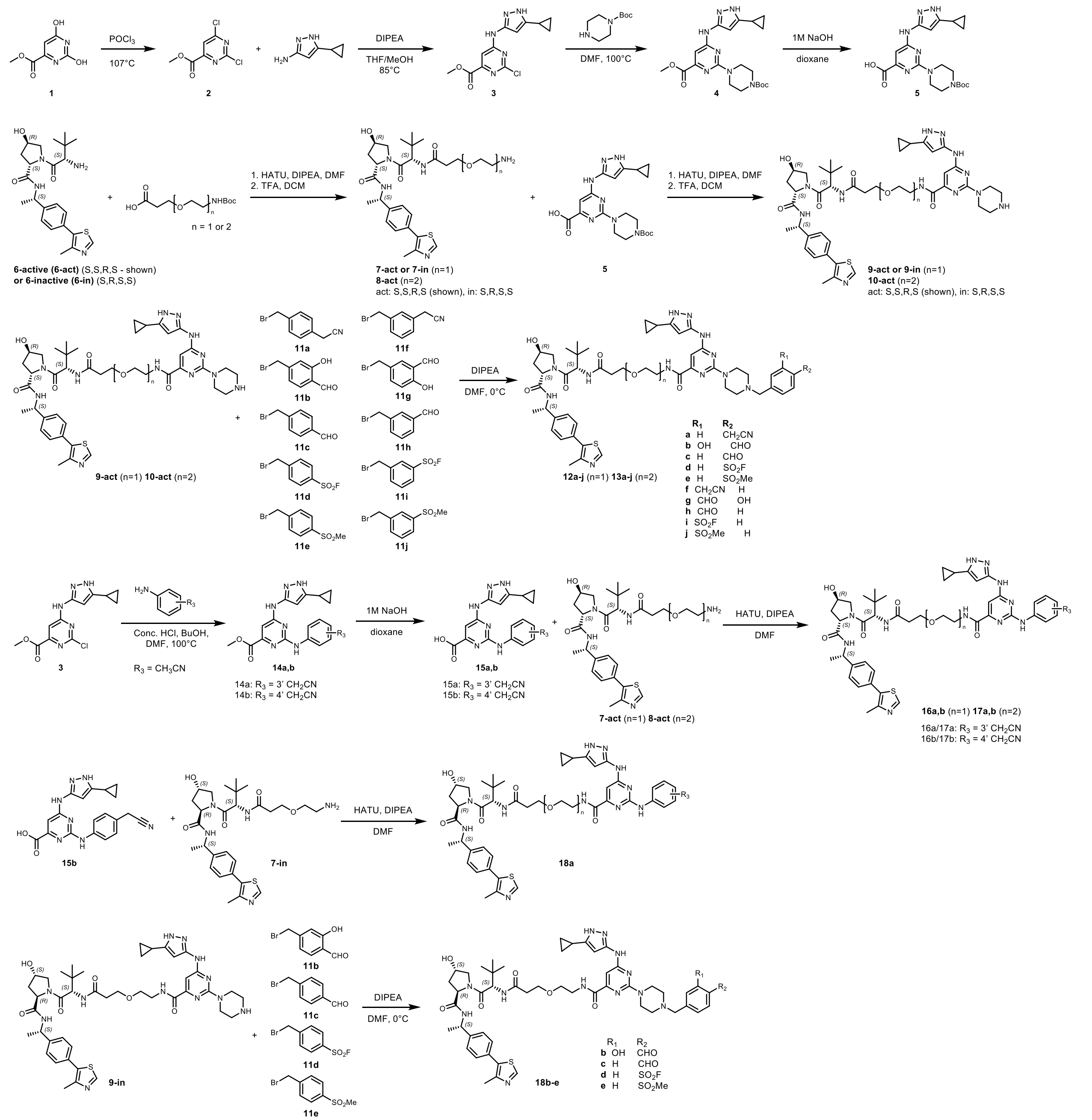

Scheme 1: Synthesis of kinetic scout degraders.

Synthetic Methods:

**General procedure A (HATU coupling):** Add amine and DIPEA to a solution of carboxylic acid and HATU in DMF. Stir for 1-16h. Dilute with ethyl acetate and wash with water and brine. Concentrate under reduced pressure.

**General procedure B (TFA Deprotection):** Add trifluoroacetic acid to a solution of Boc-protected amine in DCM. Stir for 30min and remove solvent under reduced pressure. Crude product was dried under vacuum and directly used in subsequent steps.

**General procedure C (Displacement):** Add DIPEA to a solution of amine in DMF and cool to 0°C. Add dropwise a solution of bromomethyl benzene in DMF to the reaction mixture. Warm to room temperature and stir for 1h. Crude product was purified by HPLC and lyophilized to yield the final compound.

**General procedure D (NaOH hydrolysis):** 1M Sodium hydroxide was added slowly to a solution of the methyl ester in dioxane (1mL). Stir for 30min or until reaction mixture turns completely translucent. Acidify with 1M HCl until pH 4, filter out and dry precipitate to yield carboxylic acid product.

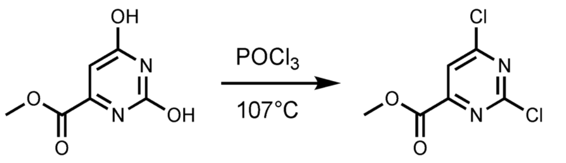


**1** **2**

**Methyl 2,6-dichloropyrimidine-4-carboxylate (2):** Suspend methyl 2,6-dihydroxypyrimidine-4-carboxylate (**1**) (3g, 0.02mol, 1eq.) in phosphorus oxychloride (30mL, 0.31mol, 15.5 eq.) at room temperature in a pressurized flask. Heat suspension to 107°C and stir for 4hrs. Dilute with DCM, transfer to a round-bottom flask, and concentrate under reduced pressure. $POCl_3$ remnants from reaction vessel and rotary evaporator was carefully poured into a large ice bath and allowed to stir overnight. Crude product was filtered and extracted with ethyl acetate. The crude product was purified using column chromatography (hexanes:ethyl acetate) to yield **2** (2.74g, 13.3mmol, 70%). LCMS $[M+H]^+$: *m/z* calcd 206.96798 found 207.1.

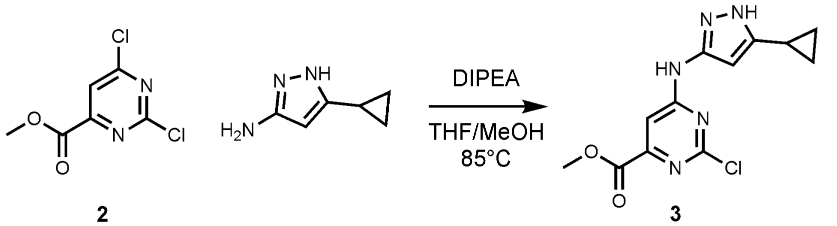


**2** **3**

**Methyl 2-chloro-6-((5-chloropropyl-1*H*-pyrazol-3-yl)amino)pyrimidine-4-carboxylate (3):** DIPEA (2.77mL, 15.9mmol, 1.2eq.) was added to a solution of 5-cyclopropyl-1*H*-pyrazol-3-amine (1.63g, 13.3mmol, 1eq.) dissolved in THF (7mL). Add a solution of **2** (2.74g, 13.3mmol, 1eq.) dissolved in THF (8mL) to the reaction mixture. Stir at room temperature for 3h. Evaporate off THF under reduced pressure and dissolve redissolve in methanol (15mL). Reflux reaction mixture at 83°C for 1h. Cool to room temperature, filter crude, and dry precipitate to yield **3** (2.72g, 9.3mmol, 70%). LCMS $[M+H]^+$: *m/z* calcd 294.06795 found 294.0.

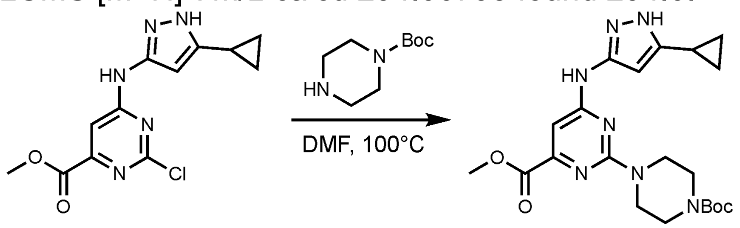


**3** **4**

**Methyl 2-(4-(*tert*-butoxycarbonyl)piperazin-1-yl)-6-((5-cyclopropyl-1*H*-pyrazol-3-yl)amino)pyrimidine-4-carboxylate (4):** *Tert*-butyl piperazine-1-carboxylate (2.3g, 13.1mmol, 2eq.) was added to a solution of **3** (1.9g, 6.5mmol, 1eq.) dissolved in DMF (50mL).The reaction mixture was heated to 100°C for 2h. Crude mixture was diluted with ethyl acetate and washed with saturated sodium bicarbonate and brine. The crude product was purified using column chromatography (hexanes:ethyl acetate) to yield **4** (1.8g, 4.12mmol, 63%). LCMS $[M+H]^+$: *m/z* calcd 390.16002 found 390.3.

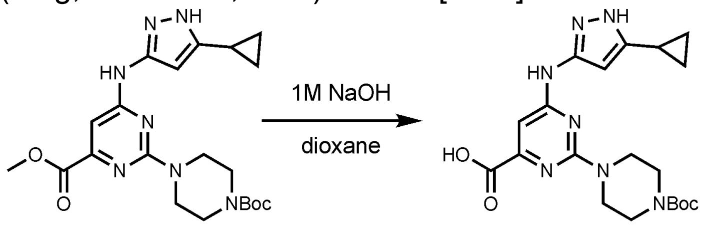


**4** **5**

**2-(4-(*tert*-butoxycarbonyl)piperazin-1-yl)-6-((5-cyclopropyl-1*H*-pyrazol-3-yl)amino)pyrimidine-4-carboxylic acid (5):** Sodium hydroxide (12mL, 1M, 3eq.), **4** (1.8g, 4.12mmol, 1eq.), and dioxane (12mL) were used according to **General Procedure D** to yield **5** (1.5g, 3.56mmol, 86%). LCMS $[M+H]^+$: *m/z* calcd 376.14437 found 376.3.

**Compound 6-active and compound 6-inactive:** Synthesized according to literature: Raina, K., *et al.* PROTAC-induced BET protein degradation as a therapy for castration-resistant prostate cancer. *PNAS* **113**, 7124–7129. https://doi.org/10.1073/pnas.1521738113

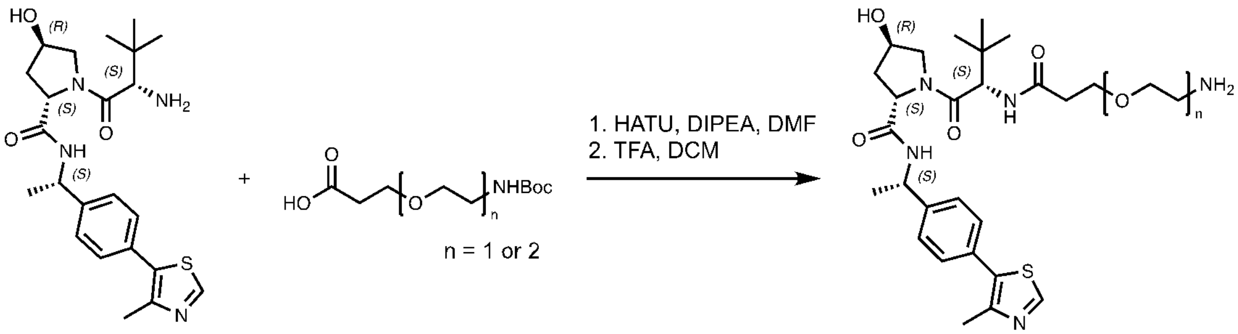


**6-active (6-act)** (S,S,R,S - shown)
**or 6-inactive (6-in)** (S,R,S,S)

**7-act or 7-in** (n=1)
**8-act** (n=2)
act: S,S,R,S (shown), in: S,R,S,S

**Compound 7-act: General procedure A** was followed using **6-act** (178mg, 0.40mmol, 1eq.), BocNH-PEG1-COOH (112mg, 0.48mmol, 1.2eq.), DIPEA (697µL, 4.0mmol, 8.3eq.), HATU (228mg, 0.60mmol, 1.5eq.), and DMF (2mL) to yield the intermediate (133mg, 0.201mmol, 50%) after purification by column chromatography (methanol:methylene chloride). LCMS $[M+H]^+$: *m/z* calcd 660.33527 found 660.5. **General procedure B** was followed using the purified intermediate (44mg, 0.066mmol, 1eq.), trifluoroacetic acid (26µL, 0.34mmol, 5eq.), and DCM (1mL) to yield **7-act** (38mg, 0.066mmol, 100%) without further purification. LCMS $[M+H]^+$: *m/z* calcd 560.28284 found 560.3.

**Compound 7-in: General procedure A** was followed using **6-in** (49mg, 0.11mmol, 1eq.), BocNH-PEG1-COOH (31mg, 0.13mmol, 1.2eq.), DIPEA (77µL, 0.44mmol, 4eq.), HATU (84mg, 0.22mmol, 2eq.), and DMF (2mL) to yield the intermediate (73mg, 0.11mmol, 100%) after purification by column chromatography (methanol:methylene chloride). LCMS $[M+H]^+$: *m/z* calcd 660.33527 found 660.6. **General procedure B** was followed using the purified intermediate (73mg, 0.11mmol, 1eq.), trifluoroacetic acid (42µL, 0.55mmol, 5eq.), and DCM (1mL) to yield **7-in** (62mg, 0.11mmol, 100%) without further purification. LCMS $[M+H]^+$: *m/z* calcd 560.28284 found 560.4.

**Compound 8-act: General procedure A** was followed using **6-act** (47.5mg, 0.11mmol, 1eq.), BocNH-PEG2-COOH (30mg, 0.11mmol, 1eq.), DIPEA (100µL, 0.57mmol, 5.3eq.), HATU (40.6mg, 0.11mmol, 1eq.), and DMF (1mL) to yield the intermediate (62.7mg, 0.089mmol, 83%) after purification by column chromatography (methanol:methylene chloride). LCMS $[M+H]^+$: *m/z* calcd 704.36148 found 704.5. **General procedure B** was followed using the purified intermediate (63mg, 0.089mmol, 1eq.), trifluoroacetic acid (34µL, 0.45mmol, 5eq.), and DCM (1mL) to yield **8-act** without further purification (54mg, 0.089mmol, 100%). LCMS $[M+H]^+$: *m/z* calcd 604.30905 found 604.4.

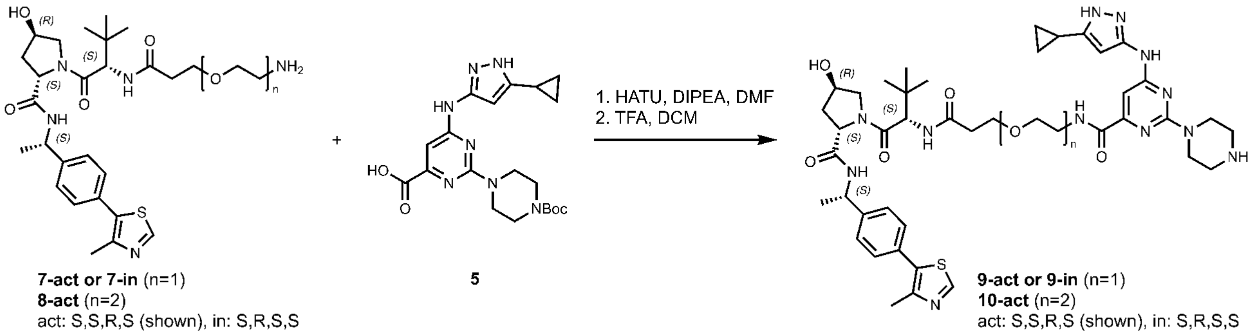


**7-act or 7-in** (n=1)
**8-act** (n=2)
act: S,S,R,S (shown), in: S,R,S,S

**5**

**9-act or 9-in** (n=1)
**10-act** (n=2)
act: S,S,R,S (shown), in: S,R,S,S

**Compound 9-act: General procedure A** was followed using the **7-act** (48mg, 0.086mmol, 1eq.), **5** (37mg, 0.086mmol, 1eq.), DIPEA (73µL, 0.43mmol, 5eq.), HATU (33mg, 0.086mmol, 1eq.), and DMF (1mL) to yield the intermediate (41mg, 0.042mmol, 49%) after purification by column chromatography (methanol:methylene chloride). LCMS $[M+H]^+$: *m/z* calcd 971.48473 found 971.9. **General procedure B** was followed using the purified intermediate (34.1mg, 0.035mmol, 1eq.), trifluoroacetic acid (13µL, 0.175mmol, 5eq.), and DCM (1ml) to yield **9-act** without further purification (31mg, 0.035mmol, 100%). LCMS $[M+H]^+$: *m/z* calcd 871.43230 found 871.9.

**Compound 9-in: General procedure A** was followed using the **7-in** (62mg, 0.11mmol, 1eq.), **5** (57mg, 0.13mmol, 1.2eq.), DIPEA (77µL, 0.44mmol, 4eq.), HATU (84mg, 0.22mmol, 2eq.), and DMF (1mL) to yield the intermediate (77mg, 0.079mmol, 72%) after purification by column chromatography (methanol:methylene chloride). LCMS $[M+H]^+$: *m/z* calcd 971.48473 found 971.8. **General procedure B** was followed using the purified intermediate (38mg, 0.039mmol, 1eq.), trifluoroacetic acid (15µL, 0.195mmol, 5eq.), and DCM (1ml) to yield **9-in** without further purification (34mg, 0.039mmol, 100%). LCMS $[M+H]^+$: *m/z* calcd 871.43230 found 871.7.

**Compound 10-act: General procedure A** was followed using the **8-act** (108mg, 0.179mmol, 1eq.), **5** (77mg, 0.179mmol, 1eq.), DIPEA (200µL, 1.0mmol, 8eq.), HATU (68mg, 0.179mmol, 1eq.), and DMF (2mL) to yield the intermediate (94mg, 0.092mmol, 51%) after purification by column chromatography (methanol:methylene chloride). LCMS $[M+H]^+$: *m/z* calcd 1015.51094 found 1015.6. **General procedure B** was followed using the purified intermediate (45mg, 0.044mmol, 1eq.), trifluoroacetic acid (17µL, 0.22mmol, 5eq.), and DCM (1ml) to yield **10-act** without further purification (41mg, 0.044mmol, 100%). LCMS $[M+H]^+$: *m/z* calcd 915.45851 found 915.7.

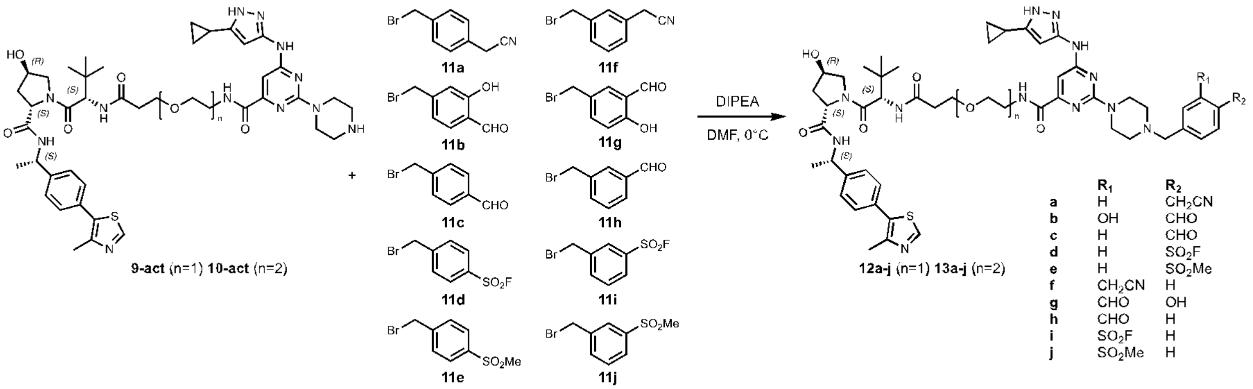


**Compound 12a (LS-1-036): General procedure C** was followed using **9-act** (5.6mg, 6.5µmol, 1eq.), **11a** (1.1mg, 6.5µmol, 1eq.), DIPEA (4.5µL, 26.1µmol, 4eq.), and DMF (1mL) to yield the final product (2.3mg, 2.1µmol, 32%) after purification by HPLC (water:methanol). $^1$H NMR (500 MHz, DMSO-$D_6$) δ 8.95 (d, *J* = 3.0 Hz, 1H), 8.58 (s, 1H), 8.36 (d, *J* = 7.8 Hz, 1H), 7.85 (d, *J* = 9.3 Hz, 1H), 7.54 (dd, *J* = 8.2, 3.0 Hz, 2H), 7.43 (d, *J* = 7.7 Hz, 2H), 7.40 (dd, *J* = 8.3, 2.9 Hz, 2H), 7.36 – 7.32 (m, 2H), 4.87 (s, 1H), 4.49 (d, *J* = 9.3 Hz, 1H), 4.39 (t, *J* = 7.9 Hz, 1H), 4.34 (s, 2H), 4.24 (s, 1H), 4.07 (d, *J* = 2.8 Hz, 2H), 3.60 – 3.54 (m, 6H), 3.44 (d, *J* = 5.8 Hz, 3H), 3.35 (s, 4H), 3.05 (s, 2H), 2.42 (d, *J* = 3.0 Hz, 3H), 2.30 (s, 1H), 1.98 (s, 1H), 1.85 (d, *J* = 9.9 Hz, 1H), 1.75 (s, 1H), 1.33 (dd, *J* = 7.0, 2.9 Hz, 3H), 1.20 (s, 1H), 0.89 (d, *J* = 2.9 Hz, 12H), 0.64 (d, *J* = 5.2 Hz, 2H). LCMS $[M+H]^+$: *m/z* calcd 1000.49015 found 1000.7.

**Compound 12b (LS-1-037): General procedure C** was followed using **9-act** (5.6mg, 6.5µmol, 1eq.), **11b** (1.4mg, 6.5µmol, 1eq.), DIPEA (4.5µL, 26.1µmol, 4eq.), and DMF (1mL) to yield the final product (3.1, 2.8µmol, 42%) after purification by HPLC (water:methanol). $^1$H NMR (500 MHz, DMSO-$D_6$) δ 11.10 (s, 1H), 10.31 (s, 1H), 8.99 (s, 1H), 8.62 (s, 1H), 8.39 (d, *J* = 7.8 Hz, 1H), 7.89 (d, *J* = 9.3 Hz, 1H), 7.75 (d, *J* = 7.9 Hz, 1H), 7.43 (d, *J* = 7.9 Hz, 2H), 7.37 (d, *J* = 7.9 Hz, 2H), 7.14 (s, 1H), 7.11 (d, *J* = 8.0 Hz, 1H), 4.94 – 4.87 (m, 1H), 4.52 (d, *J* = 9.1 Hz, 1H), 4.42 (t, *J* = 8.1 Hz, 1H), 4.36 (s, 2H), 4.27 (s, 1H), 3.60 (dd, *J* = 13.7, 8.2 Hz, 3H), 3.47 (d, *J* = 6.6 Hz, 2H), 3.39 (d, *J* = 6.9 Hz, 3H), 3.11 (s, 3H), 2.45 (s, 3H), 2.36 (d, *J* = 15.0 Hz, 1H), 2.02 (t, *J* = 10.5 Hz, 1H), 1.89 (d, *J* = 9.4 Hz, 1H), 1.78 (s, 1H), 1.36 (d, *J* = 6.9 Hz, 3H), 1.23 (s, 1H), 0.92 (s, 12H), 0.68 (d, *J* = 5.2 Hz, 2H). LCMS $[M+H]^+$: *m/z* calcd 1005.46908 found 1005.6.

**Compound 12c (JWJ-01-290): General procedure C** was followed using **9-act** (22mg, 26µmol, 1eq.), **11c** (5.6mg, 28µmol, 1.1eq.), DIPEA (6.7µL, 38.6µmol, 1.5eq.), and DMF (1mL) to yield the final product (11.3mg, 10.2µmol, 40%) after purification by HPLC (water:methanol).1H NMR (400 MHz, DMSO-D6) δ 10.04 (s, 1H), 8.95 (s, 1H), 8.58 (t, J = 6.0 Hz, 1H), 8.36 (d, J = 7.7 Hz, 1H), 8.03 – 7.95 (m, 2H), 7.86 (d, J = 9.2 Hz, 1H), 7.75 – 7.70 (m, 2H), 7.39 (d, J = 8.4 Hz, 2H), 7.33 (d, J = 8.4 Hz, 2H), 4.87 (p, J = 7.3 Hz, 1H), 4.51 – 4.42 (m, 3H), 4.38 (t, J = 8.0 Hz, 1H), 4.24 (s, 1H), 3.60 – 3.52 (m, 7H), 3.48 – 3.31 (m, 7H), 3.16 (d, J = 47.2 Hz, 4H), 2.41 (s, 3H), 2.37 – 2.27 (m, 1H), 1.98 (t, J = 10.8 Hz, 1H), 1.84 (tt, J = 8.3, 5.1 Hz, 1H), 1.74 (ddd, J = 12.9, 8.7, 4.6 Hz, 1H), 1.32 (d, J = 7.0 Hz, 3H), 0.88 (s, 12H), 0.68 – 0.59 (m, 2H). LCMS $[M+H]^+$: *m/z* calcd 989.47416 found 989.6.

**Compound 12d (JWJ-01-293): General procedure C** was followed using **9-act** (12mg, 14μmol, 1eq.), **11d** (3.8mg, 15μmol, 1.1eq.), DIPEA (3.6μL, 21μmol, 1.5eq.), and DMF (1mL) to yield the final product (7.3mg, 6.3μmol, 46%) after purification by HPLC (water:methanol). 1H NMR (400 MHz, DMSO-D6) δ 8.96 (s, 1H), 8.58 (t, J = 6.0 Hz, 1H), 8.36 (d, J = 7.8 Hz, 1H), 8.29 – 8.24 (m, 2H), 7.88 (dd, J = 12.5, 8.8 Hz, 3H), 7.42 – 7.37 (m, 2H), 7.35 – 7.31 (m, 2H), 4.87 (p, J = 7.2 Hz, 1H), 4.54 – 4.47 (m, 3H), 4.38 (t, J = 8.0 Hz, 1H), 4.24 (s, 2H), 3.63 – 3.52 (m, 5H), 3.44 (td, J = 6.2, 3.3 Hz, 3H), 3.35 (dd, J = 8.0, 4.1 Hz, 6H), 2.51 (d, J = 6.2 Hz, 1H), 2.41 (s, 3H), 2.36 – 2.27 (m, 1H), 1.98 (t, J = 11.0 Hz, 1H), 1.84 (tt, J = 8.5, 5.1 Hz, 1H), 1.32 (d, J = 6.9 Hz, 3H), 0.88 (s, 12H), 0.67 – 0.61 (m, 2H). LCMS $[M+H]^+$: *m/z* calcd 1043.43173 found 1043.4.

**Compound 12e (JWJ-01-291): General procedure C** was followed using **9-act** (12mg, 14μmol, 1eq.), **11e** (3.8mg, 15μmol, 1.1eq.), DIPEA (3.6μL, 21μmol, 1.5eq.), and DMF (1mL) to yield the final product (3.8mg, 3.3μmol, 24%) after purification by HPLC (water:methanol). 1H NMR (400 MHz, DMSO-D6) δ 8.96 (s, 1H), 8.58 (t, J = 6.0 Hz, 1H), 8.36 (d, J = 7.8 Hz, 1H), 8.05 – 7.99 (m, 2H), 7.86 (d, J = 9.3 Hz, 1H), 7.80 – 7.73 (m, 2H), 7.42 – 7.37 (m, 2H), 7.33 (d, J = 8.3 Hz, 2H), 4.87 (p, J = 7.2 Hz, 1H), 4.52 – 4.42 (m, 3H), 4.38 (t, J = 8.1 Hz, 1H), 4.24 (s, 1H), 3.58 (qd, J = 7.0, 3.3 Hz, 7H), 3.44 (tt, J = 5.7, 3.1 Hz, 3H), 3.35 (t, J = 6.1 Hz, 3H), 3.23 (s, 3H), 2.41 (s, 3H), 2.36 – 2.28 (m, 1H), 2.01 – 1.94 (m, 1H), 1.85 (tt, J = 8.4, 5.1 Hz, 1H), 1.74 (ddd, J = 12.9, 8.6, 4.5 Hz, 1H), 1.32 (d, J = 7.1 Hz, 3H), 0.88 (s, 11H), 0.67 – 0.62 (m, 2H). LCMS $[M+H]^+$: *m/z* calcd 1039.45680 found 1039.5.

**Compound 12f (LS-1-035): General procedure C** was followed using **9-act** (5.6mg, 6.5μmol, 1eq.), **11f** (1.4mg, 6.5μmol, 1eq.), DIPEA (4.5μL, 26.1μmol, 4eq.), and DMF (1mL) to yield the final product (4.2mg, 3.8μmol, 57%) after purification by HPLC (water:methanol). $^1$H NMR (500 MHz, DMSO-$D_6$) δ 10.16 (s, 1H), 8.99 (s, 1H), 8.62 (t, *J* = 6.1 Hz, 1H), 8.39 (d, *J* = 7.8 Hz, 1H), 7.89 (d, *J* = 9.3 Hz, 1H), 7.54 – 7.42 (m, 6H), 7.37 (d, *J* = 8.1 Hz, 2H), 4.91 (t, *J* = 7.2 Hz, 1H), 4.52 (d, *J* = 9.4 Hz, 1H), 4.44 – 4.38 (m, 3H), 4.28 (s, 1H), 4.12 (s, 3H), 3.61 (qd, *J* = 9.2, 3.9 Hz, 3H), 3.48 (q, *J* = 5.8 Hz, 2H), 3.45 – 3.35 (m, 3H), 3.14 (d, *J* = 26.9 Hz, 2H), 2.45 (s, 3H), 2.40 – 2.33 (m, 1H), 2.02 (t, *J* = 10.7 Hz, 1H), 1.89 (ddd, *J* = 13.5, 8.6, 5.1 Hz, 1H), 1.79 (td, *J* = 8.4, 4.2 Hz, 1H), 1.36 (d, *J* = 7.0 Hz, 3H), 1.23 (s, 1H), 0.92 (s, 12H), 0.71 – 0.64 (m, 2H). LCMS $[M+H]^+$: *m/z* calcd 1000.49015 found 1000.6.

**Compound 12g (LS-1-033): General procedure C** was followed using **9-act** (5.6mg, 6.5μmol, 1eq.), **11g** (1.3mg, 6.5μmol, 1eq.), DIPEA (4.5μL, 26.1μmol, 4eq.), and DMF

(1mL) to yield the final product (3.3mg, 2.9μmol, 45%) after purification by HPLC (water:methanol). $^{1}$H NMR (500 MHz, DMSO-$D_6$) δ 11.23 (s, 1H), 10.33 (s, 1H), 10.03 (s, 1H), 8.99 (s, 1H), 8.62 (t, *J* = 6.1 Hz, 1H), 8.39 (d, *J* = 7.8 Hz, 1H), 7.89 (d, *J* = 9.2 Hz, 1H), 7.62 (dd, *J* = 8.5, 2.3 Hz, 1H), 7.43 (d, *J* = 8.0 Hz, 2H), 7.37 (d, *J* = 8.1 Hz, 2H), 7.10 (d, *J* = 8.4 Hz, 1H), 4.91 (t, *J* = 7.2 Hz, 1H), 4.52 (d, *J* = 9.3 Hz, 1H), 4.42 (t, *J* = 8.1 Hz, 1H), 4.33 (s, 1H), 4.27 (s, 1H), 3.60 (td, *J* = 11.2, 5.9 Hz, 4H), 3.47 (q, *J* = 5.9 Hz, 2H), 3.44 – 3.37 (m, 3H), 3.26 – 3.01 (m, 3H), 2.45 (s, 3H), 2.39 – 2.33 (m, 1H), 2.02 (t, *J* = 10.8 Hz, 1H), 1.88 (td, *J* = 8.6, 4.4 Hz, 1H), 1.78 (td, *J* = 8.4, 4.2 Hz, 1H), 1.36 (d, *J* = 7.0 Hz, 3H), 1.23 (s, 1H), 0.92 (s, 12H), 0.70 – 0.66 (m, 2H). LCMS $[M+H]^+$: *m/z* calcd 1005.46908 found 1005.6.

**Compound 12h (LS-1-034): General procedure C** was followed using **9-act** (5.6mg, 6.5μmol, 1eq.), **11h** (1.4mg, 6.5μmol, 1eq.), DIPEA (4.5μL, 26.1μmol, 4eq.), and DMF (1mL) to yield the final product (2.6mg, 2.4μmol, 36%) after purification by HPLC (water:methanol). $^{1}$H NMR (500 MHz, DMSO-$D_6$) δ 10.34 (s, 1H), 10.07 (s, 1H), 8.98 (s, 1H), 8.62 (t, *J* = 6.0 Hz, 1H), 8.39 (d, *J* = 7.8 Hz, 1H), 8.08 (s, 1H), 8.04 (d, *J* = 7.6 Hz, 1H), 7.89 (d, *J* = 9.3 Hz, 1H), 7.85 (d, *J* = 7.6 Hz, 1H), 7.74 (t, *J* = 7.6 Hz, 1H), 7.43 (d, *J* = 8.0 Hz, 2H), 7.37 (d, *J* = 8.0 Hz, 2H), 4.91 (t, *J* = 7.2 Hz, 1H), 4.54 – 4.47 (m, 3H), 4.42 (t, *J* = 8.1 Hz, 1H), 4.27 (s, 1H), 3.65 – 3.56 (m, 5H), 3.47 (h, *J* = 3.6 Hz, 3H), 3.38 (d, *J* = 6.4 Hz, 3H), 3.29 – 3.09 (m, 4H), 2.45 (s, 3H), 2.40 – 2.33 (m, 1H), 2.02 (t, *J* = 10.5 Hz, 1H), 1.88 (ddd, *J* = 13.5, 8.6, 5.1 Hz, 1H), 1.78 (ddd, *J* = 13.1, 8.7, 4.7 Hz, 1H), 1.36 (d, *J* = 7.0 Hz, 3H), 1.23 (s, 1H), 0.91 (d, *J* = 4.5 Hz, 12H), 0.70 – 0.65 (m, 2H). LCMS $[M+H]^+$: *m/z* calcd 990.48199 found 989.6.

**Compound 12i (JWJ-01-294): General procedure C** was followed using **9-act** (12mg, 14μmol, 1eq.), **11i** (3.8mg, 15μmol, 1.1eq.), DIPEA (3.6μL, 21μmol, 1.5eq.), and DMF (1mL) to yield the final product (4.2mg, 4.8μmol, 38%) after purification by HPLC (water:methanol). 1H NMR (400 MHz, DMSO-D6) δ 8.95 (s, 1H), 8.59 (t, J = 6.0 Hz, 1H), 8.37 – 8.30 (m, 2H), 8.24 (dt, J = 8.0, 1.4 Hz, 1H), 8.04 (d, J = 7.7 Hz, 1H), 7.91 – 7.83 (m, 2H), 7.39 (d, J = 8.3 Hz, 2H), 7.33 (d, J = 8.4 Hz, 2H), 4.87 (p, J = 7.1 Hz, 1H), 4.54 – 4.46 (m, 3H), 4.38 (t, J = 8.0 Hz, 1H), 4.24 (s, 2H), 3.58 (ddt, J = 10.4, 7.9, 3.5 Hz, 6H), 3.48 – 3.39 (m, 4H), 3.36 (t, J = 6.0 Hz, 4H), 3.11 (d, J = 48.6 Hz, 3H), 2.50 (s, 1H), 2.41 (s, 3H), 2.35 – 2.26 (m, 1H), 1.88 – 1.67 (m, 1H), 1.32 (d, J = 7.0 Hz, 3H), 0.88 (s, 13H), 0.67 – 0.61 (m, 2H). LCMS $[M+H]^+$: *m/z* calcd 1043.43173 found 1043.5.

**Compound 12j (JWJ-01-292): General procedure C** was followed using **9-act** (12mg, 14μmol, 1eq.), **11j** (3.8mg, 15μmol, 1.1eq.), DIPEA (3.6μL, 21μmol, 1.5eq.), and DMF (1mL) to yield the final product (7.7mg, 6.7μmol, 48%) after purification by HPLC (water:methanol). 1H NMR (400 MHz, DMSO-D6) δ 8.96 (s, 1H), 8.59 (t, J = 6.0 Hz, 1H), 8.36 (d, J = 7.8 Hz, 1H), 8.09 (d, J = 2.0 Hz, 1H), 8.03 (dt, J = 7.9, 1.5 Hz, 1H), 7.90 – 7.81 (m, 2H), 7.75 (t, J = 7.7 Hz, 1H), 7.39 (d, J = 8.4 Hz, 2H), 7.33 (d, J = 8.3 Hz, 2H), 4.87 (p, J = 7.1 Hz, 2H), 4.51 – 4.43 (m, 4H), 4.38 (t, J = 8.1 Hz, 2H), 4.24 (d, J = 4.7 Hz, 3H), 3.58 (ddt, J = 10.9, 8.7, 3.7 Hz, 5H), 3.47 – 3.29 (m, 7H), 3.23 (s, 7H), 2.50 (s, 1H), 2.41 (s, 3H), 2.36 – 2.27 (m, 1H), 2.02 – 1.92 (m, 1H), 1.88 – 1.70 (m, 1H), 1.32 (d, J = 7.1 Hz, 3H), 0.88 (s, 13H), 0.68 – 0.61 (m, 2H). LCMS $[M+H]^+$: *m/z* calcd 1039.45680 found 1039.5.

**Compound 13a (LS-1-025): General procedure C** was followed using **10-act** (7.5mg, 8.2μmol, 1eq.), **11a** (1.7mg, 8.2μmol, 1eq.), DIPEA (5.7μL, 32.7μmol, 4eq.), and DMF (1mL) to yield the final product (2.0mg, 1.7μmol, 21%) after purification by HPLC (water:methanol). $^{1}$H NMR (500 MHz, DMSO-$D_6$) δ 10.22 (s, 1H), 8.99 (d, *J* = 2.6 Hz, 1H), 8.63 (d, *J* = 7.2 Hz, 1H), 8.40 (d, *J* = 7.6 Hz, 1H), 7.88 (d, *J* = 8.7 Hz, 1H), 7.56 (d, *J* = 7.9 Hz, 2H), 7.48 (d, *J* = 7.5 Hz, 2H), 7.45 – 7.41 (m, 2H), 7.37 (d, *J* = 7.7 Hz, 2H), 4.90 (t, *J* = 7.5 Hz, 1H), 4.52 (d, *J* = 9.1 Hz, 1H), 4.44 – 4.36 (m, 3H), 4.28 (s, 1H), 4.11 (d, *J* = 2.5 Hz, 3H), 3.59 (q, *J* = 10.1 Hz, 5H), 3.54 – 3.48 (m, 6H), 3.41 (s, 4H), 3.09 (s, 2H), 2.45 (d, *J* = 2.6 Hz, 3H), 2.38 – 2.30 (m, 1H), 2.01 (s, 1H), 1.92 – 1.86 (m, 1H), 1.79 (s, 1H), 1.36 (d, *J* = 7.0 Hz, 3H), 1.23 (s, 1H), 0.92 (d, *J* = 8.7 Hz, 12H), 0.68 (d, *J* = 5.4 Hz, 2H). LCMS $[M+H]^+$: *m/z* calcd 1044.51636 found 1044.9.

**Compound 13b (LS-01-032): General procedure C** was followed using **10-act** (7.5mg, 8.2μmol, 1eq.), **b**(1.8mg, 8.2μmol, 1eq.), DIPEA (5.7μL, 32.7μmol, 4eq.), and DMF (1mL) to yield the final product (4.2mg, 4.0μmol, 48%) after purification by HPLC (water:methanol). $^{1}$H NMR (500 MHz, DMSO-$D_6$) δ 11.09 (s, 1H), 10.31 (s, 1H), 9.00 (s, 1H), 8.63 (t, *J* = 6.1 Hz, 1H), 8.39 (d, *J* = 7.8 Hz, 1H), 7.87 (d, *J* = 9.3 Hz, 1H), 7.75 (d, *J* = 7.9 Hz, 1H), 7.43 (d, *J* = 8.0 Hz, 2H), 7.37 (d, *J* = 8.0 Hz, 2H), 7.14 (s, 1H), 7.10 (d, *J* = 7.9 Hz, 1H), 4.90 (t, *J* = 7.1 Hz, 1H), 4.52 (d, *J* = 9.1 Hz, 1H), 4.42 (t, *J* = 8.1 Hz, 2H), 4.37 (s, 2H), 4.27 (s, 2H), 3.59 (qd, *J* = 10.6, 4.0 Hz, 4H), 3.53 – 3.47 (m, 6H), 3.40 (d, *J* = 6.6 Hz, 3H), 3.12 (s, 3H), 2.45 (s, 3H), 2.33 (dd, *J* = 14.1, 6.7 Hz, 1H), 2.01 (t, *J* = 10.6 Hz, 1H), 1.92 – 1.86 (m, 1H), 1.78 (ddd, *J* = 12.9, 8.5, 4.7 Hz, 1H), 1.36 (d, *J* = 7.0 Hz, 3H), 1.23 (s, 1H), 0.96 – 0.88 (m, 12H), 0.68 (d, *J* = 5.2 Hz, 2H). LCMS $[M+H]^+$: *m/z* calcd 1049.49529 found 1049.5.

**Compound 13c (GJP-01-007): General procedure C** was followed using **10-act** (11.6mg, 11.3μmol, 1eq.), **11c** (2.7mg, 13.5μmol, 1.2eq.), DIPEA (2.9μL, 16.9μmol, 1.5eq.), and DMF (1mL) to yield the final product (10.2mg, 8.9μmol, 79%) after purification by HPLC (water:methanol). $^{1}$H NMR (500 MHz, DMSO-$D_6$) δ 10.25 (s, 1H), 10.08 (s, 1H), 8.99 (s, 1H), 8.62 (t, *J* = 6.2 Hz, 1H), 8.39 (d, *J* = 7.8 Hz, 1H), 8.03 (d, *J* = 7.8 Hz, 2H), 7.87 (d, *J* = 9.3 Hz, 1H), 7.76 (d, *J* = 7.8 Hz, 2H), 7.43 (d, *J* = 8.2 Hz, 2H), 7.37 (d, *J* = 8.0 Hz, 2H), 4.91 (q, *J* = 7.1 Hz, 1H), 4.54 – 4.47 (m, 3H), 4.42 (t, *J* = 8.0 Hz, 1H), 4.27 (s, 2H), 3.59 (tdd, *J* = 12.5, 9.0, 4.1 Hz, 4H), 3.54 – 3.46 (m, 6H), 3.40 (q, *J* = 6.6 Hz, 2H), 3.14 (s, 3H), 2.54 (d, *J* = 2.3 Hz, 1H), 2.45 (s, 3H), 2.37 – 2.30 (m, 1H), 2.04 – 1.98 (m, 1H), 1.88 (ddd, *J* = 13.5, 8.5, 5.0 Hz, 1H), 1.78 (ddd, *J* = 12.9, 8.6, 4.7 Hz, 1H), 1.36 (d, *J* = 7.0 Hz, 3H), 1.23 (s, 1H), 0.92 (d, *J* = 4.9 Hz, 12H), 0.72 – 0.66 (m, 2H). LCMS $[M+H]^+$: *m/z* calcd 1033.50038 found 1033.5.

**Compound 13d (AF-01-007): General procedure C** was followed using **10-act** (10mg, 11μmol, 1eq.), **11d** (2.8mg, 11μmol, 1eq.), DIPEA (7.6μL, 44μmol, 4eq.), and DMF (1mL) to yield the final product (7.9mg, 11μmol, 60%) after purification by HPLC (water:methanol). $^{1}$H NMR (500 MHz, DMSO-$D_6$) δ 8.95 (s, 1H), 8.59 (t, *J* = 6.1 Hz, 1H), 8.36 (d, *J* = 7.8 Hz, 1H), 8.26 (d, *J* = 8.2 Hz, 2H), 7.90 (d, *J* = 8.1 Hz, 2H), 7.84 (d, *J* = 9.3 Hz, 1H), 7.39 (d, *J* = 8.2 Hz, 2H), 7.33 (d, *J* = 8.1 Hz, 2H), 4.87 (t, *J* = 7.2 Hz, 1H), 4.51 (s, 1H), 4.49 (d, *J* = 9.3 Hz, 2H), 4.39 (t, *J* = 8.1 Hz, 3H), 4.24 (s, 2H), 3.56 (tt, *J* =

9.1, 4.9 Hz, 4H), 3.47 (dt, $J$ = 14.2, 5.3 Hz, 6H), 3.37 (d, $J$ = 6.4 Hz, 2H), 2.42 (s, 3H), 2.30 (dd, $J$ = 14.6, 6.3 Hz, 1H), 1.98 (t, $J$ = 10.5 Hz, 1H), 1.84 (td, $J$ = 8.5, 4.2 Hz, 1H), 1.75 (ddd, $J$ = 12.9, 8.5, 4.6 Hz, 1H), 1.32 (d, $J$ = 7.0 Hz, 3H), 0.88 (d, $J$ = 4.8 Hz, 12H), 0.67 – 0.62 (m, 2H). LCMS $[M+H]^+$: *m/z* calcd 1087.45794 found 1087.3.

**Compound 13e (AF-01-006): General procedure C** was followed using **10-act** (20mg, 22μmol, 1eq.), **11e** (5.4mg, 22μmol, 1eq.), DIPEA (15.2μL, 88μmol, 4eq.), and DMF (1mL) to yield the final product (11.7mg, 9.8μmol, 45%) after purification by HPLC (water:methanol). $^1$H NMR (500 MHz, DMSO-$D_6$) δ 10.29 (s, 1H), 10.00 (s, 1H), 9.00 (s, 1H), 8.62 (t, $J$ = 6.1 Hz, 1H), 8.39 (d, $J$ = 7.8 Hz, 1H), 8.06 (d, $J$ = 8.1 Hz, 2H), 7.87 (d, $J$ = 9.3 Hz, 1H), 7.81 (d, $J$ = 8.1 Hz, 2H), 7.43 (d, $J$ = 8.1 Hz, 2H), 7.37 (d, $J$ = 8.0 Hz, 2H), 4.90 (p, $J$ = 7.0 Hz, 1H), 4.52 (d, $J$ = 9.3 Hz, 1H), 4.50 (s, 1H), 4.42 (t, $J$ = 8.1 Hz, 1H), 4.27 (s, 1H), 3.59 (qd, $J$ = 8.1, 3.1 Hz, 4H), 3.52 – 3.49 (m, 5H), 3.41 (q, $J$ = 6.6 Hz, 3H), 3.27 (s, 3H), 3.13 (s, 1H), 2.45 (s, 3H), 2.37 – 2.30 (m, 1H), 2.05 – 1.98 (m, 1H), 1.88 (ddd, $J$ = 13.5, 8.6, 5.0 Hz, 1H), 1.78 (ddd, $J$ = 12.9, 8.6, 4.6 Hz, 1H), 1.36 (d, $J$ = 7.0 Hz, 3H), 0.92 (d, $J$ = 5.1 Hz, 11H), 0.71 – 0.65 (m, 2H). LCMS $[M+H]^+$: *m/z* calcd 1083.48301 found 1083.5.

**Compound 13f (LS-1-024): General procedure C** was followed using **10-act** (7.5mg, 8.2μmol, 1eq.), **11f** (1.7mg, 8.2μmol, 1eq.), DIPEA (5.7μL, 32.7μmol, 4eq.), and DMF (1mL) to yield the final product (2.5mg, 2.2μmol, 26%) after purification by HPLC (water:methanol). $^1$H NMR (500 MHz, DMSO-$D_6$) δ 10.17 (s, 1H), 8.95 (s, 1H), 8.60 (t, $J$ = 6.0 Hz, 1H), 8.36 (d, $J$ = 7.7 Hz, 1H), 7.84 (d, $J$ = 9.3 Hz, 1H), 7.51 – 7.46 (m, 3H), 7.43 (d, $J$ = 7.3 Hz, 1H), 7.39 (d, $J$ = 8.0 Hz, 2H), 7.33 (d, $J$ = 8.0 Hz, 2H), 4.87 (t, $J$ = 7.2 Hz, 1H), 4.49 (d, $J$ = 9.3 Hz, 1H), 4.38 (d, $J$ = 8.7 Hz, 3H), 4.24 (s, 2H), 4.08 (s, 3H), 3.55 (qd, $J$ = 10.3, 3.9 Hz, 5H), 3.49 – 3.44 (m, 6H), 3.41 – 3.35 (m, 4H), 3.06 (s, 2H), 2.42 (s, 3H), 2.29 (q, $J$ = 7.1 Hz, 1H), 1.98 (t, $J$ = 10.7 Hz, 1H), 1.85 (ddd, $J$ = 13.4, 8.6, 5.1 Hz, 1H), 1.75 (ddd, $J$ = 13.0, 8.5, 4.6 Hz, 1H), 1.33 (d, $J$ = 7.0 Hz, 3H), 1.20 (s, 1H), 0.88 (d, $J$ = 4.8 Hz, 12H), 0.65 (dd, $J$ = 5.0, 2.0 Hz, 2H). LCMS $[M+H]^+$: *m/z* calcd 1044.51636 found 1044.9.

**Compound 13g (LS-1-021): General procedure C** was followed using **10-act** (7.5mg, 8.2μmol, 1eq.), **11g** (1.4mg, 8.2μmol, 1eq.), DIPEA (5.7μL, 32.7μmol, 4eq.), and DMF (1mL) to yield the final product (1.8mg, 1.5μmol, 19%) after purification by HPLC (water:methanol). $^1$H NMR (400 MHz, DMSO-$D_6$) δ 11.22 (s, 1H), 10.33 (s, 1H), 10.00 (s, 1H), 8.99 (s, 1H), 8.64 (t, J = 6.0 Hz, 1H), 8.41 (d, J = 7.8 Hz, 1H), 7.88 (d, J = 9.3 Hz, 1H), 7.83 (d, J = 2.3 Hz, 1H), 7.62 (dd, J = 8.6, 2.4 Hz, 1H), 7.43 (d, J = 8.2 Hz, 2H), 7.37 (d, J = 8.1 Hz, 2H), 7.10 (d, J = 8.5 Hz, 1H), 4.93 – 4.87 (m, 1H), 4.52 (d, J = 9.3 Hz, 1H), 4.42 (t, J = 8.1 Hz, 1H), 4.33 (s, 2H), 4.27 (s, 2H), 3.59 (qd, J = 8.4, 3.4 Hz, 3H), 3.50 (dp, J = 11.4, 5.2 Hz, 6H), 3.44 – 3.39 (m, 3H), 3.21 (d, J = 13.6 Hz, 2H), 2.45 (s, 3H), 2.34 (dd, J = 13.4, 7.2 Hz, 1H), 2.02 (td, J = 9.3, 4.4 Hz, 1H), 1.89 (ddd, J = 13.5, 8.5, 5.1 Hz, 1H), 1.78 (td, J = 8.4, 4.3 Hz, 1H), 1.36 (d, J = 7.0 Hz, 3H), 1.23 (d, J = 4.4 Hz, 1H), 0.92 (d, J = 4.0 Hz, 12H), 0.71 – 0.66 (m, 2H).

**Compound 13h (LS-1-022): General procedure C** was followed using **10-act** (7.5mg, 8.2μmol, 1eq.), **11h** (1.6mg, 8.2μmol, 1eq.), DIPEA (5.7μL, 32.7μmol, 4eq.), and DMF

(1mL) to yield the final product (3.5mg, 3.1µmol, 37%) after purification by HPLC (water:methanol). $^1$H NMR (500 MHz, DMSO-$D_6$) δ 10.20 (s, 1H), 10.07 (s, 1H), 8.99 (s, 1H), 8.62 (t, *J* = 6.1 Hz, 1H), 8.39 (d, *J* = 7.8 Hz, 1H), 8.08 (s, 1H), 8.05 (d, *J* = 7.7 Hz, 1H), 7.90 – 7.84 (m, 2H), 7.75 (t, *J* = 7.6 Hz, 1H), 7.43 (d, *J* = 8.1 Hz, 2H), 7.37 (d, *J* = 8.1 Hz, 2H), 4.90 (t, *J* = 7.2 Hz, 1H), 4.54 – 4.47 (m, 4H), 4.42 (t, *J* = 8.1 Hz, 2H), 4.27 (s, 2H), 3.59 (tdd, *J* = 12.5, 9.4, 4.1 Hz, 4H), 3.54 – 3.48 (m, 6H), 3.41 (t, *J* = 6.3 Hz, 3H), 3.11 (d, *J* = 19.0 Hz, 4H), 2.45 (s, 3H), 2.33 (dd, *J* = 14.5, 6.3 Hz, 1H), 2.02 (d, *J* = 11.1 Hz, 1H), 1.88 (ddd, *J* = 13.5, 8.6, 5.0 Hz, 1H), 1.78 (ddd, *J* = 13.1, 8.6, 4.7 Hz, 1H), 1.36 (d, *J* = 7.0 Hz, 3H), 1.23 (s, 1H), 0.92 (d, *J* = 5.3 Hz, 12H), 0.70 – 0.66 (m, 2H). LCMS $[M+H]^+$: *m/z* calcd 1033.50038 found 1033.8.

**Compound 13i (KD-01-007): General procedure C** was followed using **10-act** (5.0mg, 5.5µmol, 1eq.), **11i** (2.8mg, 11µmol, 2eq.), DIPEA (3.8µL, 22µmol, 4eq.), and DMF (1mL) to yield the final product (4.5mg, 4.2µmol, 76%) after purification by HPLC (water:methanol). $^1$H NMR (500 MHz, DMSO-$D_6$) δ 9.96 (s, 1H), 8.96 (s, 1H), 8.60 (t, *J* = 6.1 Hz, 1H), 8.36 (d, *J* = 7.8 Hz, 1H), 8.33 (s, 1H), 8.23 (d, *J* = 8.1 Hz, 1H), 8.05 (d, *J* = 7.7 Hz, 1H), 7.88 (t, *J* = 7.9 Hz, 1H), 7.84 (d, *J* = 9.3 Hz, 1H), 7.39 (d, *J* = 8.2 Hz, 2H), 7.33 (d, *J* = 8.1 Hz, 2H), 4.87 (p, *J* = 7.0 Hz, 1H), 4.50 (s, 1H), 4.48 (s, 1H), 4.39 (t, *J* = 8.1 Hz, 1H), 4.24 (s, 1H), 3.56 (ddd, *J* = 16.1, 7.9, 4.9 Hz, 4H), 3.49 – 3.43 (m, 6H), 3.40 – 3.35 (m, 2H), 2.42 (s, 3H), 2.29 (dd, *J* = 14.6, 6.2 Hz, 1H), 2.01 – 1.96 (m, 1H), 1.85 (td, *J* = 8.6, 4.3 Hz, 1H), 1.75 (ddd, *J* = 13.0, 8.5, 4.6 Hz, 1H), 1.32 (d, *J* = 6.9 Hz, 3H), 1.20 (s, 1H), 0.89 (s, 12H), 0.67 – 0.62 (m, 2H). LCMS $[M+H]^+$: *m/z* calcd 1087.45794 found 1087.6.

**Compound 13j (KD-01-006): General procedure C** was followed using **10-act** (20mg, 22µmol, 1eq.), **11j** (5.4mg, 22µmol, 1eq.), DIPEA (12.2µL, 88µmol, 4eq.), and DMF (1mL) to yield the final product (10.6mg, 9.8µmol, 45%) after purification by HPLC (water:methanol). $^1$H NMR (500 MHz, DMSO-$D_6$) δ 10.21 (s, 1H), 8.96 (s, 1H), 8.60 (t, *J* = 6.0 Hz, 1H), 8.36 (d, *J* = 7.8 Hz, 1H), 8.10 (s, 1H), 8.03 (d, *J* = 7.9 Hz, 1H), 7.84 (d, *J* = 9.3 Hz, 2H), 7.75 (t, *J* = 7.7 Hz, 1H), 7.40 (d, *J* = 8.2 Hz, 2H), 7.33 (d, *J* = 8.1 Hz, 2H), 4.87 (t, *J* = 7.2 Hz, 1H), 4.49 (d, *J* = 8.5 Hz, 3H), 4.38 (t, *J* = 8.1 Hz, 2H), 4.24 (s, 1H), 3.61 – 3.52 (m, 4H), 3.49 – 3.43 (m, 6H), 3.38 (t, *J* = 6.3 Hz, 2H), 3.23 (s, 3H), 3.13 (s, 1H), 2.42 (s, 3H), 2.31 (dd, *J* = 14.0, 6.9 Hz, 1H), 1.98 (dd, *J* = 12.5, 8.9 Hz, 1H), 1.86 (td, *J* = 8.5, 4.3 Hz, 1H), 1.75 (ddd, *J* = 13.0, 8.6, 4.6 Hz, 1H), 1.33 (d, *J* = 7.0 Hz, 3H), 1.20 (s, 1H), 0.88 (d, *J* = 5.0 Hz, 12H), 0.67 – 0.63 (m, 2H). LCMS $[M+H]^+$: *m/z* calcd 1083.48301 found 1083.6.

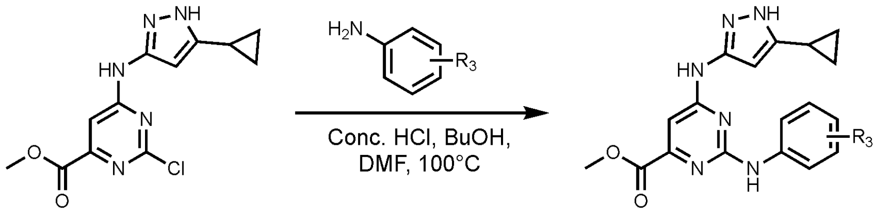


**3** $R_3 = CH_3CN$ **14a,b**

14a: $R_3$ = 3' $CH_2CN$
14b: $R_3$ = 4' $CH_2CN$

**Compound 14a:** 2-(3-aminophenyl)acetonitrile (38mg, 0.29mmol, 1.7eq.) and **3** (50mg, 0.17mmol, 1eq.) were dissolved in 2mL of butanol and 0.5mL of DMF for solubility. Concentrated HCl (28µL, 2eq.) was added to the reaction mixture and heated to 100°C overnight. Crude mixture was diluted with ethyl acetate and washed with water and brine. The crude product was purified using column chromatography (hexanes:ethyl acetate) to yield **14a** (29.9mg, 0.077mmol, 45%). LCMS $[M+H]^+$: *m/z* calcd 390.16002 found 390.3.

**Compound 14b:** 2-(4-aminophenyl)acetonitrile (27mg, 0.20mmol, 1.2eq.) and **3** (50mg, 0.17mmol, 1eq.) were dissolved in 1mL of butanol and 0.5mL of DMF for solubility. Concentrated HCl (28µL, 2eq.) was added to the reaction mixture and heated to 100°C overnight. Crude mixture was diluted with ethyl acetate and washed with water and brine. The crude product was purified using column chromatography (hexanes:ethyl acetate) to yield **14b** (42mg, 0.11mmol, 63%). LCMS $[M+H]^+$: *m/z* calcd 390.16002 found 390.2.

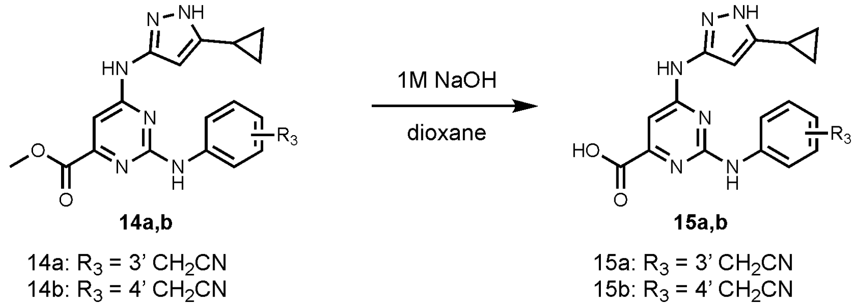


**Compound 15a:** Sodium hydroxide (0.23mL, 1M, 3eq.), **14a** (30mg, 0.077mmol, 1eq.), and dioxane (1mL) were used according to **General Procedure D** to yield **15a** (4.7mg, 0.013mmol, 16%). LCMS $[M+H]^+$: *m/z* calcd 376.14437 found 376.3.

**Compound 15b:** Sodium hydroxide (0.4mL, 1M, 3eq.), **14b** (50mg, 0.13mmol, 1eq.), and dioxane (1mL) were used according to **General Procedure D** to yield **15b** (43mg, 0.11mmol, 89%). LCMS $[M+H]^+$: *m/z* calcd 376.14437 found 376.3.

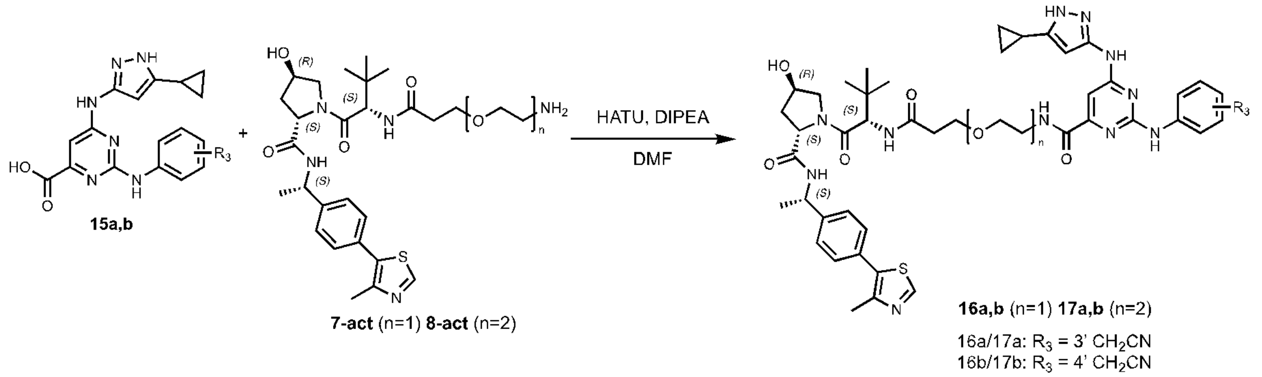


**Compound 16a (JWJ-01-289): General procedure A** was followed using **7-act** (19mg, 0.034mmol, 1eq.), **15a** (19mg, 0.051mmol, 1.5eq.), DIPEA (30µL, 0.17mmol, 5eq.), HATU (26mg, 0.068mmol, 2eq.), and DMF (1mL) to yield **16a** (5.5mg, 5.3µmol, 16%) after purification by HPLC. 1H NMR (400 MHz, DMSO-D6) δ 8.95 (s, 1H), 8.36 (d,

J = 7.8 Hz, 1H), 7.88 (d, J = 9.3 Hz, 1H), 7.71 (s, 1H), 7.62 (d, J = 8.2 Hz, 1H), 7.39 (d, J = 8.3 Hz, 2H), 7.33 (d, J = 8.4 Hz, 2H), 7.28 (t, J = 7.9 Hz, 1H), 6.92 (d, J = 7.6 Hz, 1H), 4.86 (p, J = 7.1 Hz, 1H), 4.48 (d, J = 9.4 Hz, 1H), 4.38 (t, J = 8.1 Hz, 1H), 4.23 (t, J = 3.6 Hz, 1H), 3.99 (s, 3H), 3.60 – 3.46 (m, 9H), 3.44 – 3.38 (m, 3H), 2.56 – 2.49 (m, 1H), 2.41 (s, 3H), 2.38 – 2.27 (m, 1H), 1.97 (ddd, J = 12.9, 7.6, 2.6 Hz, 1H), 1.84 (tt, J = 8.3, 5.0 Hz, 1H), 1.73 (ddd, J = 12.9, 8.6, 4.6 Hz, 1H), 1.32 (d, J = 7.1 Hz, 3H), 0.92 – 0.82 (m, 12H), 0.69 – 0.62 (m, 2H). LCMS $[M+H]^+$: *m/z* calcd 917.41665 found 917.6.

**Compound 16b (ERS-01-006): General procedure A** was followed using **7-act** (21mg, 0.037mmol, 1eq.), **15b** (14mg, 0.037mmol, 1eq.), DIPEA (32µL, 0.19mmol, 5eq.), HATU (15.3mg, 0.041mmol, 1.1eq.), and DMF (1mL) to yield **16b** (3.3mg, 3.2µmol, 9%) after purification by HPLC. $^1$H NMR (400 MHz, DMSO-$D_6$) δ 10.11 (s, 1H), 9.40 (s, 1H), 8.99 (s, 1H), 8.40 (d, *J* = 7.8 Hz, 1H), 8.21 (s, 1H), 7.93 (d, *J* = 9.3 Hz, 1H), 7.45 – 7.39 (m, 2H), 7.36 (d, *J* = 8.4 Hz, 2H), 7.27 (d, *J* = 8.2 Hz, 2H), 4.90 (p, *J* = 7.0 Hz, 1H), 4.53 (d, *J* = 9.4 Hz, 1H), 4.42 (t, *J* = 8.1 Hz, 1H), 3.97 (s, 2H), 3.46 (s, 5H), 2.60 – 2.52 (m, 1H), 2.45 (s, 3H), 2.41 – 2.36 (m, 1H), 2.01 (d, *J* = 20.4 Hz, 1H), 1.87 (t, *J* = 6.4 Hz, 1H), 1.77 (ddd, *J* = 12.9, 8.6, 4.6 Hz, 1H), 1.35 (d, *J* = 7.0 Hz, 3H), 1.23 (s, 2H), 0.91 (s, 12H), 0.72 – 0.62 (m, 3H). LCMS $[M+H]^+$: *m/z* calcd 917.41665 found 917.7.

**Compound 17a (GJP-01-008): General procedure A** was followed using **8-act** (8mg, 0.013mmol, 1eq.), **15a** (5mg, 0.013mmol, 1eq.), DIPEA (12µL, 0.69mmol, 5eq.), HATU (7mg, 0.020mmol, 1.5eq.), and DMF (1mL) to yield **17a** (3.9mg, 4.1µmol, 30%) after purification by HPLC. $^1$H NMR (400 MHz, DMSO-$D_6$) δ 10.22 (s, 1H), 9.50 (s, 1H), 8.99 (s, 1H), 8.40 (d, *J* = 7.8 Hz, 1H), 8.24 (s, 1H), 7.87 (d, *J* = 9.3 Hz, 1H), 7.74 (s, 1H), 7.65 (d, *J* = 8.4 Hz, 1H), 7.43 (d, *J* = 8.3 Hz, 2H), 7.37 (d, *J* = 8.3 Hz, 2H), 7.32 (t, *J* = 7.9 Hz, 1H), 6.96 (d, *J* = 7.6 Hz, 1H), 4.91 (p, *J* = 7.0 Hz, 1H), 4.52 (d, *J* = 9.4 Hz, 1H), 4.42 (t, *J* = 8.1 Hz, 2H), 4.27 (d, *J* = 3.9 Hz, 2H), 4.03 (s, 3H), 3.58 (t, *J* = 4.9 Hz, 2H), 3.55 – 3.52 (m, 4H), 3.48 (ddt, *J* = 8.3, 4.8, 3.0 Hz, 4H), 2.45 (s, 3H), 2.37 – 2.28 (m, 1H), 2.01 (dd, *J* = 12.9, 8.1 Hz, 1H), 1.88 (tt, *J* = 8.4, 5.1 Hz, 1H), 1.77 (ddd, *J* = 12.9, 8.6, 4.6 Hz, 1H), 1.36 (d, *J* = 7.0 Hz, 3H), 1.27 – 1.20 (m, 1H), 0.92 (s, 12H), 0.73 – 0.67 (m, 2H). LCMS $[M+H]^+$: *m/z* calcd 961.44286 found 961.5.

**Compound 17b (GJP-01-004): General procedure A** was followed using **8-act** (33mg, 0.055mmol, 1eq.), **15b** (21mg, 0.055mmol, 1eq.), DIPEA (48µL, 0.27mmol, 5eq.), HATU (22.6mg, 0.060mmol, 1.1eq.), and DMF (1mL) to yield **17b** (12.6mg, 11.7mmol, 21%) after purification by HPLC. $^1$H NMR (500 MHz, DMSO-$D_6$) δ 10.29 (s, 1H), 9.46 (s, 1H), 8.99 (s, 1H), 8.39 (d, *J* = 7.8 Hz, 1H), 8.34 (s, 1H), 7.87 (d, *J* = 9.3 Hz, 1H), 7.69 (d, *J* = 8.2 Hz, 2H), 7.47 – 7.40 (m, 2H), 7.37 (d, *J* = 8.2 Hz, 2H), 7.29 (d, *J* = 8.2 Hz, 2H), 4.91 (q, *J* = 7.2 Hz, 1H), 4.52 (d, *J* = 9.4 Hz, 2H), 4.42 (t, *J* = 8.1 Hz, 3H), 4.27 (s, 2H), 3.99 (s, 2H), 3.63 – 3.57 (m, 4H), 3.57 – 3.53 (m, 5H), 3.52 – 3.43 (m, 4H), 2.45 (s, 3H), 2.37 – 2.32 (m, 1H), 2.01 (t, *J* = 9.8 Hz, 1H), 1.87 (tt, *J* = 8.6, 4.9 Hz, 1H), 1.78 (ddd, *J* = 12.9, 8.6, 4.7 Hz, 1H), 1.36 (d, *J* = 7.0 Hz, 3H), 1.23 (s, 1H), 0.94 – 0.89 (m, 12H), 0.69 – 0.65 (m, 2H). LCMS $[M+H]^+$: *m/z* calcd 961.44286 found 961.6.

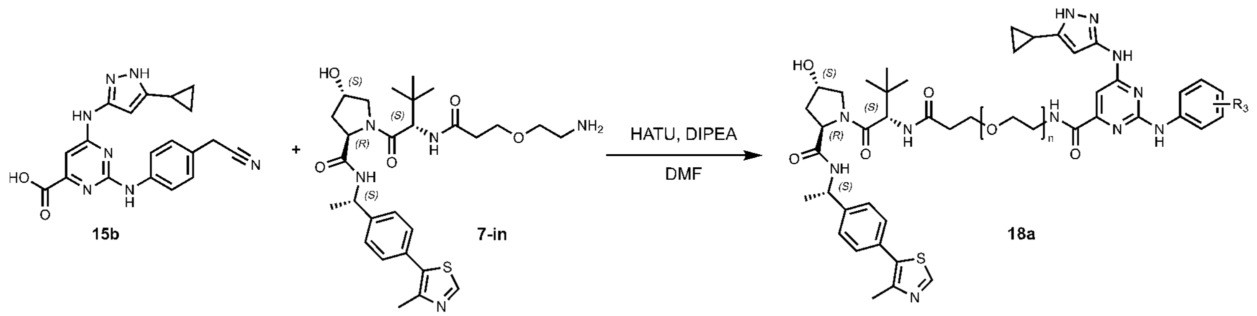


**Compound 18a (ATF-01-129): General procedure A** was followed using **7-in** (20mg, 0.036mmol, 1eq.), **15b** (15mg, 0.039mmol, 1.1eq.), DIPEA (25µL, 0.14mmol, 4eq.), HATU (27.2mg, 0.072mmol, 2eq.), and DMF (1mL) to yield **16b** (3.5mg, 3.4µmol, 10%) after purification by HPLC. $^1$H NMR LCMS $[M+H]^+$: *m/z* calcd 917.41665 found 917.8. $^1$H NMR (400 MHz, DMSO-D6) δ 9.43 (s, 1H), 8.96 (d, J = 3.6 Hz, 1H), 8.27 (s, 1H), 8.09 (d, J = 8.0 Hz, 1H), 8.03 (d, J = 8.1 Hz, 1H), 7.68 (d, J = 7.9 Hz, 2H), 7.42 (s, 4H), 7.27 (d, J = 8.3 Hz, 2H), 4.88 (p, J = 7.1 Hz, 1H), 4.43 (d, J = 8.1 Hz, 1H), 4.39 – 4.35 (m, 1H), 4.30 (q, J = 4.8 Hz, 1H), 3.97 (s, 2H), 3.76 (dd, J = 10.5, 5.2 Hz, 1H), 3.70 – 3.62 (m, 1H), 3.59 (dt, J = 9.7, 6.5 Hz, 1H), 3.48 (dt, J = 11.9, 5.1 Hz, 3H), 3.41 (p, J = 5.4 Hz, 2H), 2.61 – 2.53 (m, 1H), 2.44 (s, 3H), 2.37 – 2.29 (m, 1H), 2.07 – 1.96 (m, 1H), 1.96 – 1.90 (m, 1H), 1.89 – 1.83 (m, 1H), 1.30 (d, J = 7.0 Hz, 3H), 0.93 (s, 12H), 0.67 (s, 2H).

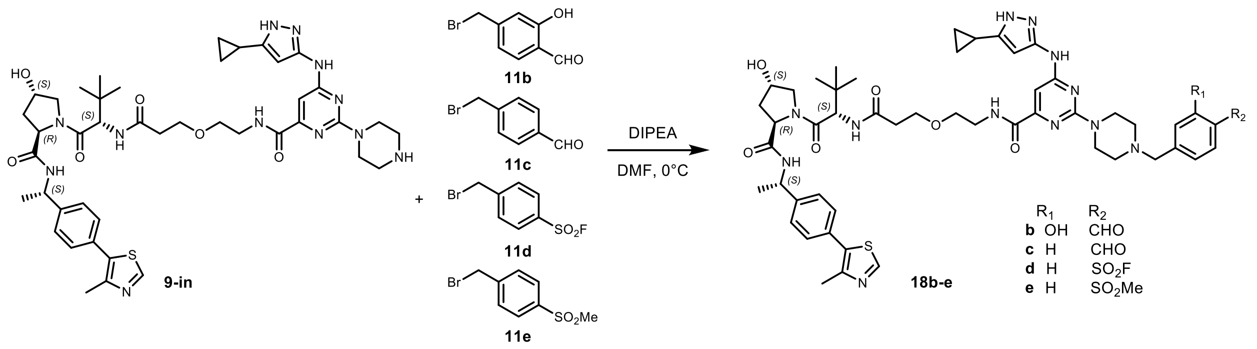


**Compound 18b (ATF-01-076): General procedure C** was followed using **9-in** (20mg, 23µmol, 1eq.), **11b** (5.5mg, 28µmol, 1.2eq.), DIPEA (9.8µL, 56µmol, 2eq.), and DMF (1mL) to yield the final product (4.9mg, 4.4µmol, 19%) after purification by HPLC (water:methanol). $^1$H NMR (500 MHz, DMSO-$D_6$) δ 11.06 (s, 1H), 10.31 (s, 1H), 8.97 (s, 1H), 8.59 (s, 1H), 8.10 (d, *J* = 8.0 Hz, 1H), 7.99 (d, *J* = 8.2 Hz, 1H), 7.75 (d, *J* = 7.9 Hz, 1H), 7.54 – 7.46 (m, 1H), 7.42 (s, 4H), 7.13 (s, 1H), 7.10 (d, *J* = 7.9 Hz, 1H), 4.88 (t, *J* = 7.4 Hz, 1H), 4.43 (d, *J* = 8.1 Hz, 1H), 4.40 – 4.34 (m, 3H), 4.31 (t, *J* = 5.0 Hz, 1H), 3.62 – 3.56 (m, 4H), 3.53 – 3.48 (m, 2H), 3.43 (t, *J* = 5.8 Hz, 3H), 3.11 (s, 2H), 2.59 – 2.53 (m, 1H), 2.44 (s, 3H), 2.31 (dd, *J* = 13.9, 6.8 Hz, 1H), 1.99 (t, *J* = 6.7 Hz, 1H), 1.95 – 1.86 (m, 2H), 1.31 (d, *J* = 6.9 Hz, 3H), 0.94 (s, 12H), 0.68 (q, *J* = 3.9 Hz, 3H). LCMS $[M+H]^+$: *m/z* calcd 1005.46908 found 1005.7.

**Compound 18c (ATF-01-074): General procedure C** was followed using **9-in** (20mg, 23µmol, 1eq.), **11c** (5.9mg, 28µmol, 1.2eq.), DIPEA (9.8µL, 56µmol, 2eq.), and DMF (1mL) to yield the final product (3.9mg, 3.5µmol, 15%) after purification by HPLC (water:methanol). $^1$H NMR (500 MHz, DMSO-$D_6$) δ 10.08 (s, 1H), 8.97 (s, 1H), 8.59 (t, *J* = 6.0 Hz, 1H), 8.10 (d, *J* = 8.0 Hz, 1H), 8.05 – 8.00 (m, 2H), 7.99 (d, *J* = 8.2 Hz, 1H), 7.75 (d, *J* = 7.9 Hz, 2H), 7.42 (d, *J* = 2.1 Hz, 4H), 4.88 (t, *J* = 7.4 Hz, 1H), 4.48 (s, 2H), 4.43 (d, *J* = 8.1 Hz, 1H), 4.38 (dd, *J* = 8.2, 5.9 Hz, 1H), 4.32 (d, *J* = 5.2 Hz, 1H), 3.77 (dd, *J* = 10.7, 5.2 Hz, 2H), 3.63 – 3.55 (m, 2H), 3.50 (d, *J* = 10.8 Hz, 1H), 3.43 (q, *J* = 5.8 Hz, 3H), 3.34 (d, *J* = 7.3 Hz, 2H), 2.44 (d, *J* = 2.0 Hz, 3H), 2.32 (dt, *J* = 13.3, 6.1 Hz, 1H), 1.99 (t, *J* = 6.7 Hz, 1H), 1.93 (t, *J* = 5.9 Hz, 1H), 1.91 – 1.85 (m, 1H), 1.30 (d, *J* = 7.0 Hz, 3H), 1.00 – 0.85 (m, 12H), 0.67 (d, *J* = 5.8 Hz, 3H). LCMS $[M+H]^+$: *m/z* calcd 989.47416 found 989.8.

**Compound 18d (ATF-01-075): General procedure C** was followed using **9-in** (20mg, 23μmol, 1eq.), **11d** (7.0mg, 28μmol, 1.2eq.), DIPEA (9.8μL, 56μmol, 2eq.), and DMF (1mL) to yield the final product (6.8mg, 5.9μmol, 26%) after purification by HPLC (water:methanol). $^1$H NMR (500 MHz, DMSO-$D_6$) δ 8.97 (d, *J* = 2.2 Hz, 1H), 8.59 (d, *J* = 6.3 Hz, 1H), 8.31 – 8.27 (m, 2H), 8.10 (d, *J* = 8.0 Hz, 1H), 7.99 (d, *J* = 8.2 Hz, 1H), 7.92 (d, *J* = 8.1 Hz, 2H), 7.42 (d, *J* = 2.2 Hz, 4H), 4.89 (q, *J* = 7.3 Hz, 1H), 4.53 (s, 2H), 4.43 (d, *J* = 8.3 Hz, 1H), 4.38 (dd, *J* = 8.2, 5.9 Hz, 1H), 4.32 (d, *J* = 6.1 Hz, 2H), 3.77 (dd, *J* = 10.5, 5.1 Hz, 2H), 3.65 – 3.55 (m, 2H), 3.50 (d, *J* = 10.3 Hz, 1H), 3.43 (q, *J* = 5.7 Hz, 3H), 2.56 (dd, *J* = 13.8, 6.6 Hz, 1H), 2.44 (d, *J* = 2.1 Hz, 3H), 2.36 – 2.28 (m, 1H), 1.99 (t, *J* = 6.7 Hz, 1H), 1.94 (t, *J* = 5.9 Hz, 1H), 1.88 (ddd, *J* = 13.5, 9.0, 5.5 Hz, 1H), 1.31 (d, *J* = 6.9 Hz, 3H), 0.98 – 0.88 (m, 12H), 0.68 (q, *J* = 3.5 Hz, 3H). LCMS $[M+H]^+$: *m/z* calcd 1043.43173 found 1043.6.

**Compound 18e (ATF-01-162): General procedure C** was followed using **9-in** (20mg, 23μmol, 1eq.), **11e** (6.9mg, 28μmol, 1.2eq.), DIPEA (9.8μL. 56μmol, 2eq.), and DMF (1mL) to yield the final product (4.4mg. 3.8μmol, 17%) after purification by HPLC (water:methanol). $^1$H NMR (400 MHz, DMSO-$D_6$) δ 8.97 (s, 1H), 8.61 (t, *J* = 6.0 Hz, 1H), 8.15 – 8.09 (m, 1H), 8.09 – 8.04 (m, 2H), 8.01 (d, *J* = 8.2 Hz, 1H), 7.81 – 7.77 (m, 2H), 7.42 (s, 4H), 4.88 (p, *J* = 7.2 Hz, 1H), 4.49 (s, 2H), 4.43 (d, *J* = 8.2 Hz, 1H), 4.38 (dd, *J* = 8.1, 5.9 Hz, 1H), 4.32 (q, *J* = 4.9 Hz, 1H), 3.65 – 3.53 (m, 1H), 3.50 (dd, *J* = 10.5, 3.8 Hz, 1H), 3.43 (td, *J* = 6.0, 3.6 Hz, 2H), 3.34 (q, *J* = 6.7 Hz, 2H), 3.26 (s, 3H), 2.44 (s, 3H), 2.31 (dt, *J* = 14.4, 6.0 Hz, 1H), 2.01 (t, *J* = 6.4 Hz, 1H), 1.98 – 1.91 (m, 1H), 1.91 – 1.85 (m, 1H), 1.30 (d, *J* = 7.0 Hz, 3H), 0.94 (s, 9H), 0.68 (dt, *J* = 3.4, 2.2 Hz, 2H). LCMS $[M+H]^+$: *m/z* calcd 1039.45680 found 1039.8.
