## Supplementary material for "A Kinetic Scout Approach Accelerates Targeted Protein Degrader Development": Biology methods

**Cell culture**

The following cell lines were employed in this study: HEK293 (ATCC, CRL-1573), HEK293FT-FKBP12F36VNluc (N. S. Gray lab), MOLT4 (ATCC,CVCL_0013LT), and MOLT4 VHL-/- (N. S. Gray lab). HEK293 and HEK293FT-FKBP12F36VNluc were cultured in high-glucose DMEM (Gibco, 11965092) with 10% FBS (Gibco, 10437028) and 1% Penicillin–Streptomycin (Gibco, 15140122). MOLT4 and MOLT4 VHL-/- were cultured with RPMI-1640 media (Gibco, 11875093) with 10% FBS (Gibco, 10437028) and 1% Penicillin–Streptomycin (Gibco, 15140122). All cell lines were maintained in 37 °C and 5% CO2 incubators and routinely tested negative for mycoplasma contamination using the MycoAlert Kit (Lonza, LT07318).

**NanoBRET VHL Cellular Target Engagement Assay**

The Promega VHL NanoBRET TE Assay (Promega, N2931) was used to quantify VHL target engagement. HEK293 cells were transfected with transfection mixture containing 9.0 μg/ml Transfection Carrier DNA, 1.0 μg/ml of VHL-NanoLuc® fusion vector DNA and TransIT-2020 (Mirus, 5400) in Opti-MEM (Gibco, 11058021). The transfection complex was allowed to incubate for 20 min. HEK293 cells were obtained and adjusted to a density of 200,000 cells/mL in Opti-MEM with 1% FBS and combined with transfection complex at a 20:1 cell to transfection complex ratio. The cells were transferred into a tissue culture flask and incubated for 24 hr at 37°C, 5% CO2. The following day, 34 μL of cells at 200,000 cells/mL in Opti-MEM were plated into 384-well white, nonbinding surface microplates (Corning, 3574) and co-treated in triplicate with 1 μM tracer and a 10-point half-log titration of kinetic scout degraders starting at 100 μM in DMSO using a Multidrop Pico 8 Digital Dispenser (Thermo). The plates were then incubated at 37°C, 5% CO2 for 2 hr. Plates were allowed to equilibrate to room temperature for 15 min prior to characterization. Then 17 μL of 3X Complete Substrate Plus Inhibitor Solution was added to each well of the 384-well plate. Plates were incubated for 2 min at room temperature and then donor emission wavelength (450nm) and acceptor emission wavelength (610nm) were measured using a ClarioSTAR Plus microplate reader (BMG Labtech). To calculate BRET ratios the acceptor signal/emission signal for the no-tracer background control was subtracted from the acceptor signal/emission signal for the treatment condition and then that value was multiplied by 1000. These values were then normalized to the DMSO BRET ratios. Data was plotted in GraphPad Prism and EC50 calculated using a log(inhibitor) vs response – variable slope.

**dTAG Competition VHL Target Engagement Assay**

Target engagement assay was adapted from Nabet et al. using the Nano-Glo Dual-Luciferase Reporter Assay System (Promega, N1630). Briefly, HEK293FT-FKBP12F36V-NanoLuc cells were plated in 20 μL of culture media at 4,000 cells per well in a TC-treated 384-well white microplate (Corning, 3570). Cells were allowed to adhere to plates overnight in 37°C, 5% CO2 incubator and the following day the cells were co-treated in triplicate with 100 nM dTAG-v and a 10-point half-log titration of kinetic scout degraders starting at 100 μM in DMSO using a Multidrop Pico 8 Digital Dispenser (Thermo). Cells were incubated for 6 hr at 37°C, 5% CO2 and then brought to room temperature 15 min prior to characterization. Cells were treated with 20 μL of ONE-Glo EX reagent and incubated on an orbital shaker for 10 min at 600 rpm then Firefly luminesce was measured using a ClarioSTAR Plus microplate reader (BMG Labtech). Afterward, cells were treated with 20 μL of NanoDLR Stop and Glo reagent and incubated on an orbital shaker for 10 min at 600 rpm, then NanoLuc luminescence measured. To normalize the data, the NanoLuc values were divided by the Firefly values and the resulting

NanoLuc/Firefly ratios were normalized by the respective DMSO ratios. Data was plotted in GraphPad Prism and EC50 calculated using a log(inhibitor) vs response – variable slope.

#### Cell Viability Assay

MOLT4WT and MOLT4VHL-/- cells were cultured and plated into TC-treated white 384-well microplate (Corning, 3570) at 800 cells in 20 µL culture media per well, then returned to the 37°C, 5% $CO_2$ incubator overnight. Cells were treated in triplicate with DMSO or kinetic scout degrader in a 10-point half-log titration starting at 100 µM using a Multidrop Pico 8 Digital Dispenser (Thermo), then returned to incubator for 72 hr. Cells were equilibrated to room temperature, then 10 µL reconstituted CellTiter-Glo (Promega, G7571) reagent was added to each well. Microplate was mixed for 2 min at 600 rpm on an orbital shaker then incubated for 10 min at room temperature. Luminescence was measured using a plate reader (CLARIOstar Plus, BMG Labtech). Raw data was DMSO normalized, log-transformed, and plotted in Prism (GraphPad). EC50 was calculated using the built-in "[Inhibitor] vs. response – Variable slope (four parameters)" analysis method.

#### Western Blot

MOLT4 cells were plated into a 12-well plate (Corning, 3513) at 3,000,000 cells per well in culture media. Each well was treated with indicated concentration of kinetic scout degrader and incubated for 6 hr. Cells were then harvested and pellets were washed with 500 µL PBS and then resuspended in 50 µL M-PER lysis buffer (Thermo, 78503) containing 1X HALT inhibitor cocktail (Thermo, 78442). Tubes were then lysed on ice for 1 hr. Lysates were clarified by centrifuging at 21,000 x g for 15 min at 4°C. Protein concentration was determined by bicinchoninic acid (BCA) assay (Thermo, 23225). SDS-PAGE samples were prepared in 1X NuPage LDS sample buffer (Invitrogen, NP0007) and boiled at 95 °C for 5 min. SDS-PAGE samples were run on a Bolt 4–12% Bis-Tris Gel (Invitrogen, NW04125BOX) in MES run buffer (Invitrogen, B0002) for 45 min at 180 V, then transferred to a nitrocellulose membrane (Cytiva, 10600011) in Bolt transfer buffer (Invitrogen, BT00061) for 90 min at 45 V. Membrane was blocked with 5% non-fat dry milk (Kroger) in TBST (Thermo, 28360) for 1 hr at room temperature, then incubated with primary antibody diluted in TBS blocking buffer (LI-COR, 92760001) overnight at 4 °C. Primary antibodies used were 1:1000 CDK6 (Cell Signaling Technology, 13331T), 1:1000 NEK9 (Abcam, ab138488) and 1:1000 alpha-tubulin (Cell Signaling Technology, 3873S). Membrane was washed three times with TBST then incubated with DyLight 680 anti-mouse IgG (Cell Signaling Technology, 5470S) and DyLight 800 anti-rabbit IgG (Cell Signaling Technology, 5151S) diluted 1:10000 in TBS blocking buffer for 1 hr at room temperature. Membrane was washed three times with TBST then imaged with a ChemiDoc Imaging System (Bio-Rad) and quantified in Image Lab 6.1.0 (Bio-Rad).

#### CDK6 Target Engagement Assay

The Promega K-10 NanoBRET TE Intracellular Kinase Assay (Promega, N2640) was used to quantify CDK6 target engagement. HEK293 cells were transfected with transfection mixture containing 9.0 µg/ml Transfection Carrier DNA, 1.0 µg/ml of CDK6-NanoLuc® fusion vector DNA and TransIT-2020 (Mirus, 5400) in Opti-MEM (Gibco, 11058021). The transfection complex was allowed to incubate for 20 min. HEK293 cells were obtained and adjusted to a density of 200,000 cells/mL in Opti-MEM with 1% FBS and combined with transfection complex at a 20:1 cell to transfection complex ratio. Then 34 µL of cells were plated into 384-well white microplate (Corning, 3570). Cells were allowed to incubate at 37°C, 5% CO2 for 24 hr and the following day the cells were co-treated in triplicate with 0.5 µM K-10 tracer and a 10-point half-log titration of negative control kinetic scout degraders starting at 100 µM in DMSO using a Multidrop Pico 8 Digital Dispenser (Thermo). The plates were then incubated at 37°C, 5% CO2 for 2 hr. Plates were allowed to equilibrate to room temperature for 15 min prior to characterization. Then 17 µL of 3X Complete Substrate Plus Inhibitor Solution was added to each well of the 384-well plate. Plates were incubated for 2 min at room temperature and then donor emission wavelength (450nm) and acceptor emission wavelength (610nm) were measured using a ClarioSTAR Plus microplate reader (BMG Labtech). To calculate BRET ratios the acceptor signal/emission signal for the no-tracer background control was subtracted from the acceptor signal/emission signal for the treatment condition and then that value was multiplied by 1000. These values were then normalized to the DMSO BRET ratios. Data was plotted in GraphPad Prism and EC50 calculated using a log(inhibitor) vs response – variable slope.

**CDK6 Washout Target Engagement Assay**

The Promega K-10 NanoBRET TE Intracellular Kinase Assay (Promega, N2640) was used to quantify CDK6 target engagement after compound washout. HEK293 cells were transfected with transfection mixture containing 9.0 µg/ml Transfection Carrier DNA, 1.0 µg/ml of CDK6-NanoLuc® fusion vector DNA and TransIT-2020 (Mirus, 5400) in Opti-MEM (Gibco, 11058021). The transfection complex was allowed to incubate for 20 min. HEK293 cells were obtained and adjusted to a density of 200,000 cells/mL in Opti-MEM and combined with transfection complex at a 20:1 cell to transfection complex ratio. Then 85 µL of cells were plated into 96-well white microplate (Corning, 3917). Cells were allowed to incubate at 37°C, 5% CO2 for 24 hr and the following day the cells were with 10 µM negative control kinetic scout degraders using a Multidrop Pico 8 Digital Dispenser (Thermo). The plates were then incubated at 37°C, 5% CO2 for 2 hr. The media was removed from the wells intended to be washed. Twice, 85 µL of Opti-MEM containing 10% FBS was added to each well and incubated for 5 minutes and then removed from the well. Following the FBS containing washes, two additional washes were performed using 85 µL of Opti-MEM. After the washes, the wells were treated with 0.5 µM K-10 tracer. Immediately after the addition of the tracer, 42.5 µL of 3X Complete Substrate Plus Inhibitor Solution was added to each well of the 96-well plate. Plates were incubated for 2 min at room temperature and then donor emission wavelength (450nm) and acceptor emission wavelength (610nm) were measured using a ClarioSTAR Plus microplate reader (BMG Labtech). Measurements were taken every 3 minutes for 120 minutes. To calculate BRET ratios the acceptor signal/emission signal for the no-tracer background control was subtracted from the acceptor signal/emission signal for the treatment condition and then that value was multiplied by 1000. Data was plotted in GraphPad Prism.

**K192 Target Engagement Assay**

The Promega NanoBRET K192 Kinase Selectivity System (Promega, NP4050) was used to quantify target engagement across the kinome following a similar protocol described in Nieman et al. Briefly, HEK293 cells were cultivated and split one day before transfection to yield cells that are 80% confluent on transfection day. NanoLuc-Kinase vector plates were resuspended using sterile TE buffer to give 20 ng/µL DNA. To perform the transfections, 8 µL of Opti-MEM

(Gibco, 11058021) was added to each well of a 384-well white microplate (Corning, 3570) and then 2 µL of each DNA was transferred to its respective well of the 384-well plate. TransIT-2020 (Mirus, 5400) was diluted in Opti-MEM at a ratio of 0.16 µL TransIT-2020 to 5 µL Opti-MEM and 5 µL diluted TransIT-2020 was added to each transfection well. The transfection complexes were allowed to form for 30 min. 5 µL of HEK293 cells in Opti-MEM + 4% FBS (Gibco, 10437028) was added to each well to give 7,000 cells per well. Cells were allowed to incubate at 37°C, 5% CO2 for 24 hr and the following day the cells were co-treated in triplicate with K-10 tracer (at concentration indicated by kit) and 1 µM negative control kinetic scout degrader in DMSO using a Multidrop Pico 8 Digital Dispenser (Thermo). The plates were then incubated at 37°C, 5% CO2 for 2 hr. Plates were allowed to equilibrate to room temperature for 15 min prior to characterization. Then 10 µL of 3X Complete Substrate Plus Inhibitor Solution was added to each well of the 384-well plate. Plates were incubated for 2 min at room temperature and then donor emission wavelength (450nm) and acceptor emission wavelength (610nm) were measured using a ClarioSTAR Plus microplate reader (BMG Labtech). To calculate BRET ratio values the acceptor signal was divided by the emission signal for each well. To calculate the percent target engagement the BRET ratio value for a given condition was divided by its respective DMSO BRET ratio value and this value was then subtracted from 1 and multiplied by 100.

### Global quantitative proteomics

Cells were lysed by addition of lysis buffer (8 M Urea, 50 mM NaCl, 50 mM 4-(2-hydroxyethyl)-1-piperazineethanesulfonic acid (EPPS) pH 8.5, Protease and Phosphatase inhibitors) and homogenization by bead beating (BioSpec) for three repeats of 30 seconds at 2400 strokes/min. Bradford assay was used to determine the final protein concentration in the clarified cell lysate. Fifty micrograms of protein for each sample was reduced, alkylated and precipitated using methanol/chloroform as previously described [1] and the resulting washed precipitated protein was allowed to air dry. Precipitated protein was resuspended in 4 M urea, 50 mM HEPES pH 7.4, followed by dilution to 1 M urea with the addition of 200 mM EPPS, pH 8. Proteins were digested with the addition of LysC (1:50; enzyme:protein) and trypsin (1:50; enzyme:protein) for 12 h at 37 °C. Sample digests were acidified with formic acid to a pH of 2-3 before desalting using C18 solid phase extraction plates (SOLA, Thermo Fisher Scientific). Desalted peptides were dried in a vacuum-centrifuged and reconstituted in 0.1% formic acid for liquid chromatography-mass spectrometry analysis.

Data were collected using a TimsTOF Pro2 (Bruker Daltonics, Bremen, Germany) coupled to a nanoElute LC pump (Bruker Daltonics, Bremen, Germany) via a CaptiveSpray nano-electrospray source. Peptides were separated on a reversed-phase C18 column (25 cm x 75 µM ID, 1.6 µM, IonOpticks, Australia) containing an integrated captive spray emitter. Peptides were separated using a 50 min gradient of 2 - 30% buffer B (acetonitrile in 0.1% formic acid) with a flow rate of 250 nL/min and column temperature maintained at 50 ºC.

To perform diaPASEF, the precursor distribution in the DDA m/z-ion mobility plane was used to design an acquisition scheme for DIA data collection which included two windows in each 50 ms diaPASEF scan. Data was acquired using sixteen of these 25 Da precursor double window scans (creating 32 windows) which covered the diagonal scan line for doubly and triply charged precursors, with singly charged precursors able to be excluded by their position in the m/z-ion mobility plane. These precursor isolation windows were defined between 400 - 1200 m/z and 1/k0 of 0.7 - 1.3 V.s/cm2.

The diaPASEF raw file processing and controlling peptide and protein level false discovery rates, assembling proteins from peptides, and protein quantification from peptides was performed using library free analysis in DIA-NN 1.8[83]. Library free mode performs an in-silico digestion of a given protein sequence database alongside deep learning-based predictions to extract the DIA precursor data into a collection of MS2 spectra. The search results are then used to generate a spectral library which is then employed for the targeted analysis of the DIA data searched against a Swissprot human database (January 2021). Database search criteria largely followed the default settings for directDIA including: tryptic with two missed cleavages, carbamidomethylation of cysteine, and oxidation of methionine and precursor Q-value (FDR) cut-off of 0.01. Precursor quantification strategy was set to Robust LC (high accuracy) with RT-dependent cross run normalization.

Resulting data was filtered to only include proteins that had a minimum of 2 counts in at least 4 replicates of each independent comparison of treatment sample to the DMSO control. Proteins with missing values were imputed by random selection from a Gaussian distribution either with a mean of the non-missing values for that treatment group or with a mean equal to the median of the background (in cases when all values for a treatment group are missing). Significant changes comparing the relative protein abundance of these treatment to DMSO control comparisons were assessed by moderated t test as implemented in the limma package within the R framework (M.E. Ritchie et al., 2015, Nucleic Acids Res, 43(7):e47).

**E-STUB sample processing and quantitative proteomics**

HEK293T cells stably expressing VHL-BirA fusion proteins were cultured in DMEM media supplemented with 10% biotin-depleted FBS and transfected with pRK5-A3-HA-ubiquitin. One day after transfection, cells were treated with carfilzomib (0.4 µM) (Selleck Chemicals, S2853) for an hour, followed by DMSO or drug treatment for 1 hour, during which the last 15 minutes was a biotin pulse (50 µM). Three biological replicates were prepared for each treatment condition. Cell harvesting and sample preparation for analysis by mass spectrometry were performed exactly as previously described (Huang *et al*, Nat. Chem. Biol., 2024).

Data were collected using a TimsTOF Pro2 (Bruker Daltonics, Bremen, Germany) coupled to a nanoElute LC pump (Bruker Daltonics, Bremen, Germany) via a CaptiveSpray nano-electrospray source. Peptides were separated on a reversed-phase C18 column (25 cm x 75 µM ID, 1.6 µM, IonOpticks, Australia) containing an integrated captive spray emitter. Peptides were separated using a 50 min gradient of 2 - 30% buffer B (acetonitrile in 0.1% formic acid) with a flow rate of 250 nL/min and column temperature maintained at 50 ºC.

To perform diaPASEF, the precursor distribution in the DDA m/z-ion mobility plane was used to design an acquisition scheme for DIA data collection which included two windows in each 50 ms diaPASEF scan. Data was acquired using sixteen of these 25 Da precursor double window scans (creating 32 windows) which covered the diagonal scan line for doubly and triply charged precursors, with singly charged precursors able to be excluded by their position in the m/z-ion mobility plane. These precursor isolation windows were defined between 400 - 1200 m/z and 1/k0 of 0.7 - 1.3 V.s/cm2.

The diaPASEF raw file processing and controlling peptide and protein level false discovery rates, assembling proteins from peptides, and protein quantification from peptides was performed using library free analysis in DIA-NN 1.8[83]. Library free mode performs an in-silico digestion of a given protein sequence database alongside deep learning-based predictions to extract the DIA precursor data into a collection of MS2 spectra. The search results are then used to generate a spectral library which is then employed for the targeted analysis of the DIA data searched against a Swissprot human database (January 2021). Database search criteria largely followed the default settings for directDIA including: tryptic with two missed cleavages, carbamidomethylation of cysteine, and oxidation of methionine and precursor Q-value (FDR) cut-off of 0.01. Precursor quantification strategy was set to Robust LC (high accuracy) with RT-dependent cross run normalization.

Resulting data was filtered to only include proteins that had a minimum of 2 counts in at least 2 replicates of each independent comparison of treatment sample to the DMSO control. Protein abundances were globally normalized using in-house scripts in the R framework (R Development Core Team, 2014). Proteins with missing values were imputed by random selection from a Gaussian distribution either with a mean of the non-missing values for that treatment group or with a mean equal to the median of the background (in cases when all values for a treatment group are missing). Significant changes comparing the relative protein abundance of these treatment to DMSO control comparisons were assessed by moderated t test as implemented in the limma package within the R framework (M.E. Ritchie et al., 2015, Nucleic Acids Res, 43(7):e47).

**Data Availability**

Raw proteomics data have been deposited to the PRIDE repository hosted by ProteomeXchange, with the following PXD archive numbers.

Global proteomics: PXD055400 (wp-esf_458)

VHL IP/MS: PXD055398 (ip-esf_360)

E-STUB: PXD055399 (ip-esf_381)
